# The power of universal contextualised protein embeddings in cross-species protein function prediction

**DOI:** 10.1101/2021.04.19.440461

**Authors:** Irene van den Bent, Stavros Makrodimitris, Marcel Reinders

## Abstract

Computationally annotating proteins with a molecular function is a difficult problem that is made even harder due to the limited amount of available labelled protein training data. A recently published supervised molecular function predicting model partly circumvents this limitation by making its predictions based on the universal (i.e. task-agnostic) contextualised protein embeddings from the deep pre-trained unsupervised protein language model SeqVec. SeqVec embeddings incorporate contextual information of amino acids, thereby modelling the underlying principles of protein sequences insensitive to the context of species.

We applied the existing SeqVec-based molecular function prediction model in a transfer learning task by training the model on annotated protein sequences of one training species and making predictions on the proteins of several test species with varying evolutionary distance. We show that this approach successfully generalises knowledge about protein function from one eukaryotic species to various other species, proving itself an effective method for molecular function prediction in inadequately annotated species from understudied taxonomic kingdoms. Furthermore, we submitted the performance of our SeqVec-based prediction models to detailed characterisation, first to advance the understanding of protein language models and second to determine areas of improvement.

**Author summary:** Proteins are diverse molecules that regulate all processes in biology. The field of synthetic biology aims to understand these protein functions to solve problems in medicine, manufacturing, and agriculture. Unfortunately, for many proteins only their amino acid sequence is known whereas their function remains unknown. Only a few species have been well-studied such as mouse, human and yeast. Hence, we need to increase knowledge on protein functions. Doing so is, however, complicated as determining protein functions experimentally is time-consuming, expensive, and technically limited. Computationally predicting protein functions offers a faster and more scalable approach but is hampered as it requires much data to design accurate function prediction algorithms. Here, we show that it is possible to computationally generalize knowledge on protein function from one well-studied training species to another test species. Additionally, we show that the quality of these protein function predictions depends on how structurally similar the proteins are between the species. Advantageously, the predictors require only the annotations of proteins from the training species and mere amino acid sequences of test species which may particularly benefit the function prediction of species from understudied taxonomic kingdoms such as the Plantae, Protozoa and Chromista.

## Introduction

Proteins are diverse molecules that perform many different functions in cells ranging from catalyzing chemical reactions to functioning as mere structural components [1–3]. These functions are generally described in terms of functional Gene Ontology (GO) annotations. GO annotations, also known as GO terms, are statements about the molecular function of the protein, cellular localisation of the protein or the biological process it supports [4]. This knowledge on protein function has come to play a central role in our daily lives, fueling the field of synthetic biology and thereby solving problems in medicine, manufacturing and agriculture [5–7].

To date, however, most GO annotations linked to proteins are shallow and incomplete [8, 9]. Additionally, as increasingly more protein sequences are characterised by high-throughput wet-lab experiments, they often remain without any functional annotation [10, 11]. Especially in certain taxonomic kingdoms such as the Plantae, Protozoa and Chromista, very few species have been thoroughly studied and the quality of available annotations is substandard [12].

Extensive wet-lab experiments remain the most accurate tools to annotate proteins but are time-consuming, expensive and some proteins cannot be studied at all due to technical limitations [13]. In response, there have been numerous attempts to functionally annotate proteins using automated, fast and scalable bioinformatics tools [14, 15]. Early approaches like BLAST often rely on homology relationships to identify conserved protein sequences, transferring the functional annotation between them as conserved sequence implies conserved function [16, 17]. These kinds of approaches quickly drop in predictive power for divergent proteins, and therefore new generation approaches often integrate numerous types of protein data with potent computational tools like neural networks. These types of protein data include sequence motifs, structural motifs, co-expression data and protein-protein interactions usually extracted from (the combination of) amino acid sequences, 3D structures and high-throughput techniques [14, 18, 19]. Whereas the new generation techniques usually outperform the established BLAST baseline method, they also require vast amounts of protein data which is not always comprehensive. Therefore, the most recent approaches often turn to automatic representation learning by which a complex model (often a neural network) learns some abstract features of a protein sequence which contain useful information for a consequent computational function prediction task [20–22].

Recently, we demonstrated that features generated from the pre-trained protein language model Seqvec [23] significantly outperform methods that learn sequence features in a supervised manner even when coupled with a simple linear classifier [22]. The advantage of language models is that they are trained in an unsupervised fashion, by training to predict each amino acid in a protein sequence given its ‘context’, i.e. neighboring amino acids [23–26]. This means that they can be trained on all available protein sequences and not only the annotated ones. By leveraging this wealth of data, they can learn general properties of amino acids, such as polarity and secondary structure, that are very useful for downstream prediction tasks [23], while deep supervised methods can be limited by the relatively small number of proteins available for training [22].

More specifically, language models learn a fixed-length vector for each amino acid, often referred to as an amino acid embedding. High-quality embeddings should ideally model both (i) complex characteristics of the side chains (e.g. size, polarity, charge), and (ii) how the effects of these side chains vary across environmental context (i.e. model side-chain interactions with spatially close side chains). We refer to this as (i) the ‘semantics’ of amino acids, and (ii) the ‘syntax’ or ‘underlying principles’ of protein sequences. To represent an entire protein by automatic representation learning, a protein-level embedding is usually obtained from amino acid-level embeddings, effectively summarizing the information contained in them. Another advantage of protein-level embeddings in function prediction tasks lies in their simplicity; after the model that produces the embeddings is trained, protein embeddings are easily obtained as they do not require sequence alignments, structural data, or selection of informative amino acid properties.

The SeqVec model is based on the ELMo model from the field of Natural Language Processing (NLP) [27] and consists of three layers: an amino acid embedding layer (aaEM), a first bidirectional Long Short-Term Memory (biLSTM 1) and a second biLSTM (biLSTM 2) layer (Figure 1A). It is trained to predict the next or previous amino acid in the protein sequence given all the previous or following amino acids. To this end, it uses an independent forward and backward pass in the biLSTM layers, respectively. During inference each layer of the SeqVec model produces a 1024-dimensional embedding for every amino acid in the protein sequence (Figure 1B). As a standard approach introduced by [23], these three embeddings are summed component-wise to obtain a final amino acid-level embedding. Consequently, SeqVec protein-level embeddings are obtained by calculating the component-wise mean over the sequence of amino acid-level embeddings (Figure 1C). This results in a final 1024-dimensional protein-level embedding independent of the protein sequence length.

**Fig 1.**
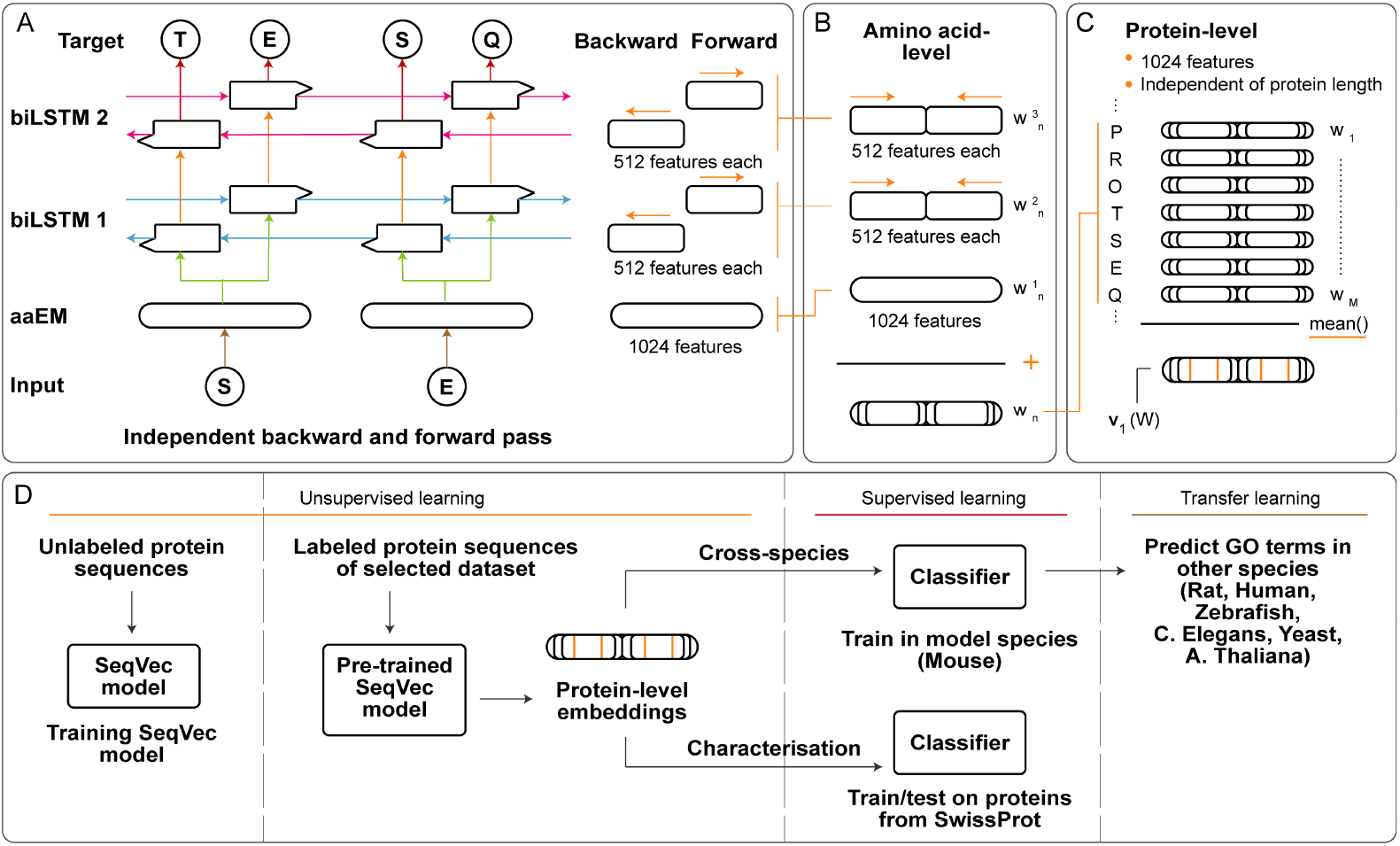
(A) SeqVec model architecture holds 3 layers: an amino acid embedding layer (aaEM), the first bidirectional Long Short Term Memory (biLSMT 1) and the second biLSTM (biLSTM 2). The aaEM maps the input amino acid onto a latent 1024-dimensional space. Both biLSTM’s have a separate forward and backward pass to incorporate information on the previous or following amino acids and map this information onto 512-dimensional spaces. Based on the biLSMT 2 embeddings, the model predicts the next or previous amino acid in the sequence. Weights between the forward and backward pass are shared, represented here as arrows with the same colour. (B) During inference, the embeddings of both biLSTM layers are concatenated to 1024-dimensional embeddings. Using the standard approach from [23], a final amino acid-level embedding is obtained by summing the 1024-dimensional embeddings **w^1^**_*n*_, **w^2^**_*n*_ and **w^3^**_*n*_ resulting in a 1024-dimensional contextualised embedding *w_n_* for every amino acid *n* in the protein sequence. (C) Protein-level embeddings are obtained by calculating the component-wise arithmetic mean of the sequence of amino acid embeddings (**w**_1_,…, **w**_*n*_) resulting in 1024-dimensional protein-level embedding **v**_1_ (**W**) independent of protein length. (D) Overview of approaches taken in this study. The pre-trained SeqVec model is used to embed proteins in a 1024-dimensional space. In the cross-species experiments, we train a molecular function predictor on a ‘central’ species and evaluate its performance on proteins of other species. In our characterisation experiments, we use the embeddings to get a deeper understanding of SeqVec-based molecular function prediction performance.

In this paper, we build upon previous work [22,23] and use the SeqVec embeddings to examine the reliability of predicting functions between proteins of evolutionary distant species. Till date, making predictions over large evolutionary distances is difficult as the function of a protein is determined by the context of its species [15]. If SeqVec can truly learn the underlying principles of protein sequences (i.e. the semantics and syntax of amino acids), we expect SeqVec-based protein function prediction to be much less sensitive to this limitation as SeqVec will produce universal contextualized protein-level embeddings independent of the context of species. In support of our approach, SeqVec embeddings were already shown to model the ‘semantics’ of amino acids as they model their biochemical and biophysical properties [23]. Additionally, as the SeqVec embeddings are contextualized as a result of the the biLSTM layers in the SeqVec model, they may also model the ‘syntax’ of proteins. If proven effective, this particular cross-species approach could be especially useful for understudied evolutionary kingdoms (e.g. Plantae) to readily generalise knowledge on protein function.

The concept of cross-species prediction has been previously touched upon when Jensen et al. showed that knowledge on protein ‘cellular roles’, which are much broader statements about protein functions than molecular functions, can be generalized from one eukaryotic species to other eukaryotic species [28]. They made their predictions using the ProtFun method which uses as input hand-crafted features such as post-translational and localization aspects of the protein. We extend this work by (i) using the more powerful SeqVec model and (ii) predicting more specific molecular functions instead of broad cellular roles of proteins. We train a SeqVec-based molecular function prediction model on the annotated proteins from one training species (Mouse) and assess its performance on the proteins of other, evolutionary related and distant species (Figure 1D). As proof of principle, we use the data from well-annotated eukaryotic species to aid performance evaluation which typically relies on comparing predicted functions to the true functions of proteins.

In summary, we demonstrate the effectiveness and reliability of this data-undemanding approach by successfully transferring knowledge on protein function between the training species and various other eukaryotic species. Thereby, we present an innovative method for molecular function prediction in inadequately annotated species from understudied taxonomic kingdoms.

As language models are relatively novel in the field of protein function prediction, we also submit the performance of SeqVec-based molecular function prediction models to detailed characterisation to advance the understanding of such models (Figure 1D). In so doing, we uncover a clear relationship between performance and structural conservation.

## Results characterisation

### Evaluating SeqVec-based molecular function prediction performance

We already showed that SeqVec embeddings achieve competitive performance when applied in the task of molecular function prediction [22]. We built upon this previous work to characterise SeqVec-based molecular function prediction. To this end, we used the same so-called SwissProt dataset (30% maximum-pairwise sequence similarity, 3.530 test proteins and 441 GO terms). Although our previous work revealed that a multilayer perceptron (MLP) trained on protein-level embeddings had the best performance, the MLP was also trained in a multilabel fashion. Hence, underlying relations between GO annotations and their abundance in the training set might have influenced the MLP performance in ways we are unsure about. As a Logistic Regression (LR) can be easily be trained independently for every GO term and showed only slightly lower performance than the MLP in our previous work, we use a LR to characterize the performance while minimizing influences on performance beyond our control. Throughout this study, we evaluated the performance of a classifier in a term-centric (ROCAUC score) and protein-centric (F1 score) way as proposed and described in detail by [15].

We observed similar performance values as our previous work with an average ROCAUC score of 0.832 (95% confidence interval [0.827 - 0.837]) and an average F1 score 0.479 (95% confidence interval [0.470 - 0.484]). Additionally, the coverage was 0.998, indicating that for almost all proteins in the test set at least one molecular function was predicted.

### Term-centric performance correlates positively with GO term depth and negatively with the number of training proteins

To find GO term characteristics indicative of performance we characterised the term-centric performance of each GO term to its depth and amount of training proteins. Depth is defined as the length of the longest possible path to a GO term from the root term in the GO hierarchy. We note that as a result of the GO hierarchy the depth and number of training proteins were not independent parameters, as terms closer to the root (i.e. with low depth) usually have more annotated proteins (Spearman correlation: −0.34, p-value: 1.55e-13).

We observed a weak non-linear positive correlation between GO term depth and term-centric performance (Spearman correlation: 0.16, p-value: 1.1e-3) (Figure 2A). The number of proteins in the training set and term-centric performance showed a weak non-linear negative correlation (Spearman correlation: −0.19, p-value: 6.6e-5) (Figure 2B). However, even between terms with the same depth or number of training proteins, we observed a large spread in the term-centric performance. This indicates that some molecular functions are somehow easier to predict than others. Additionally, it indicates that these differences in term-centric performance could also have roots in protein characteristics, making some proteins easier to predict than others.

**Fig 2.**
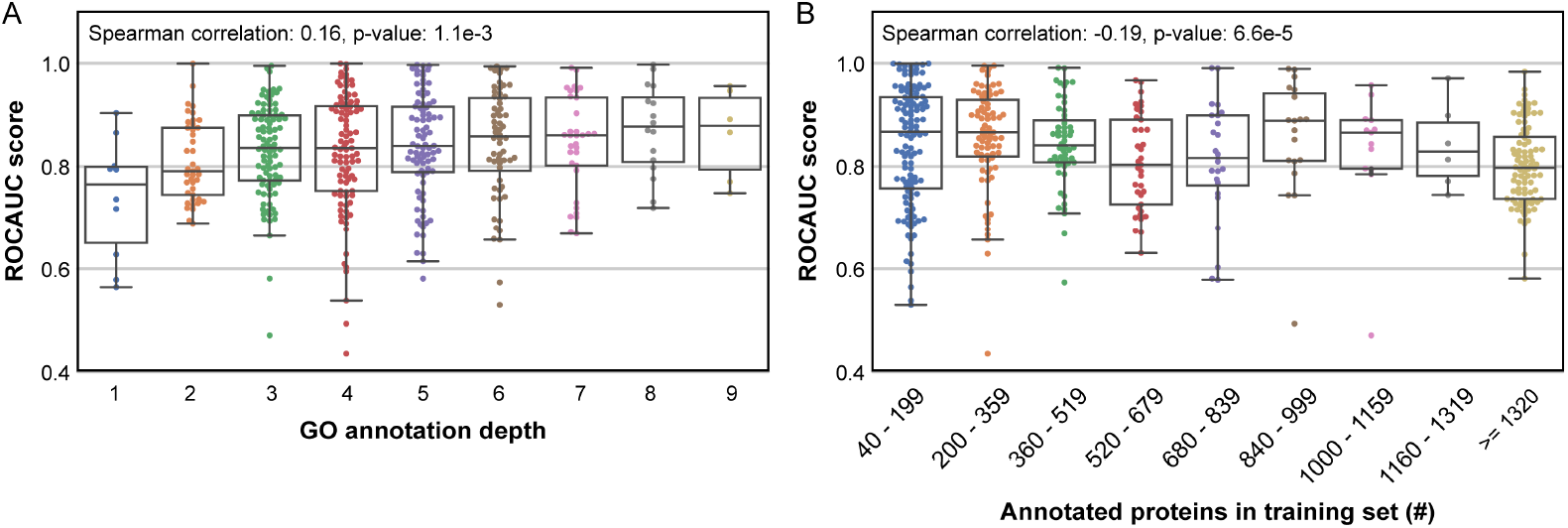
Term-centric performance (ROCAUC) per GO term of a LR classifier trained using SeqVec protein-level embeddings on the SwissProt dataset with at most 30% pairwise sequence identity in relation to (A) depth of the GO term and (B) the number of annotated proteins in the training set for the GO term. The Spearman correlations and corresponding p-values are shown in the respective plots. The box-whiskers plots show the interquartile range (IQR) with a box and the median as a bar across the box. Whiskers denote the range equal to 1.5 times the IQR.

### Protein-centric performance correlates positively with protein length

To test whether protein characteristics could be underlying the differences in term-centric performance, we characterised the protein-centric performance to protein length and the number of protein annotations. Again, these parameters were not independent as in eukaryotes the protein size is positively correlated with more extended multifunctional proteins [29, 30]. However, in the SwissProt dataset, we observed only a mild correlation which we attributed to the lack of true annotations for many proteins (Spearman correlation: 0.14, p-value: 1.55e-13) [8, 9].

We observed a weak non-linear positive correlation between protein length and protein-centric performance (Spearman coefficient: 0.10, p-value: 7.2e-9) (Figure 3A). The slightly increased protein-centric performance for longer proteins was mainly the result of improved precision for longer proteins as recall remained similar for all protein lengths (Figure S.1A, B). For an increased number of protein annotations we observed a slightly decreased protein-centric performance, although testing for a correlation resulted in no statistical significant finding (Spearman coefficient: 0.03, p-value: 0.06) (Figure 3B). Here, whereas precision increased for more annotations, recall decreased (Figure S.1C, D).

**Fig 3.**
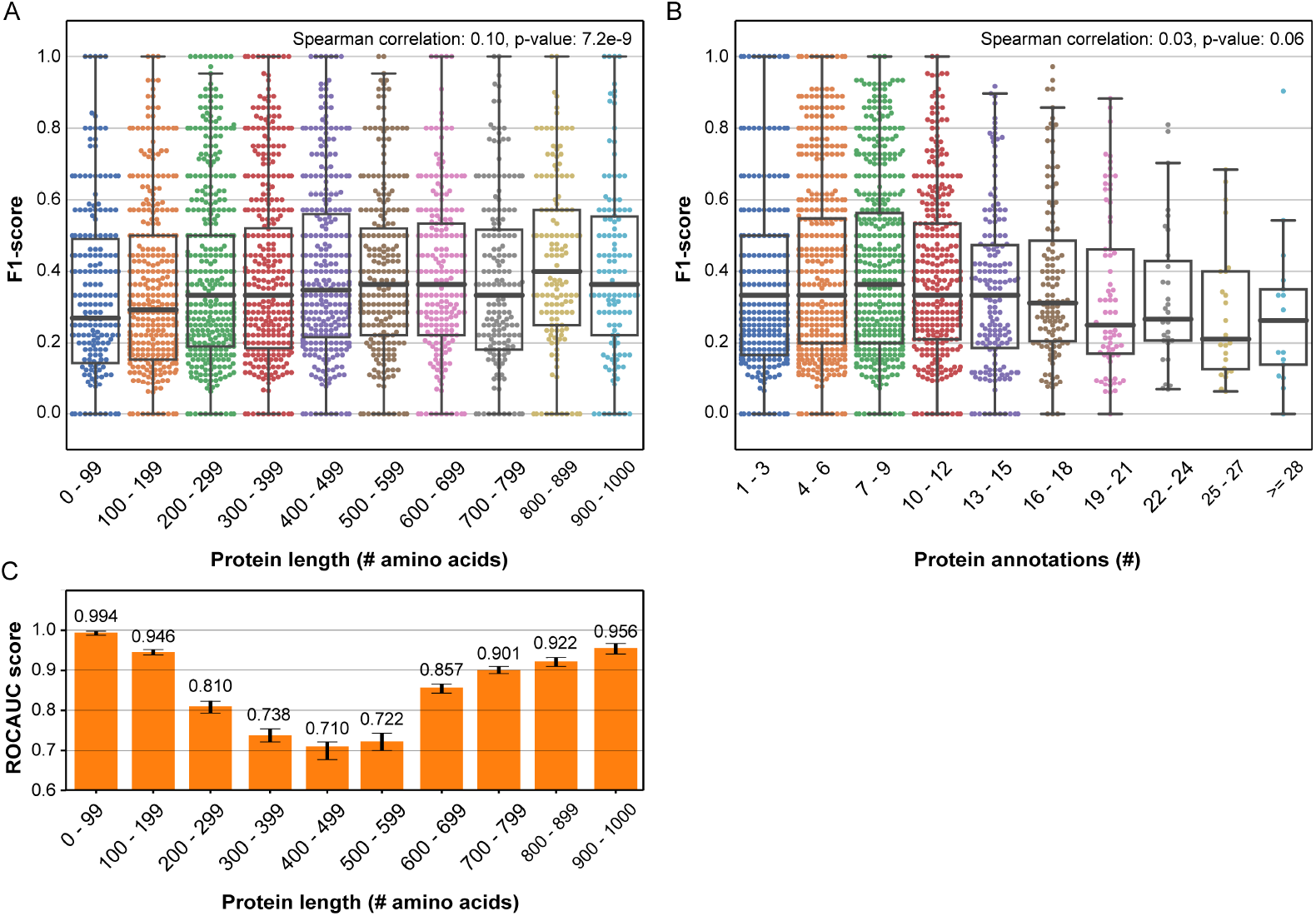
Protein-centric performance (F1) per protein of the LR classifier trained using baseline SeqVec protein-level embeddings on the SwissProt dataset in relation to (A) protein sequence length and (B) the number of protein annotations. The LR was trained to predict GO terms. The Spearman correlations and corresponding p-values between protein-centric performance and protein length or number of protein annotations are shown. The box-whiskers plots show the interquartile range (IQR) with a box and the median as a bar across the box. Whiskers denote the range equal to 1.5 times the IQR. (C) Term-centric performance (ROCAUC) of the LR classifier trained using baseline SeqVec protein-level embeddings on the SwissProt dataset. The LR was trained to predict protein length encoded by one-hot encoding in the same intervals as in (A). Errorbars denote 95% confidence estimated using 100 bootstraps.

Overall, the difference in protein-centric performance between long and short proteins was small. Therefore, even if some GO terms with the same depth or number of training proteins had large differences in the average length of their annotated proteins, our evaluated protein characteristics seemed an unlikely source of the differences in term-centric performance. However, in theory, taking the mean over a larger number of amino acid-level embeddings should discard more information and yield an average closer to the population mean, whereas taking the mean over less amino acid-level embeddings should give an average closer to the sample mean [31]. Hence, we expected a lower protein-centric performance for longer proteins. This was not the case, hinting that SeqVec embeddings likely model some protein characteristic that countered the expected decrease in performance for longer proteins.

### Protein-level embeddings effectively model protein length

To explain the observed positive correlation, we hypothesised that the protein length is somehow encoded in SeqVec embeddings. Specifically, as long proteins are relatively scarce, they might be easier to predict by having similar embeddings as other long protein in the training set with similar function. To test our hypothesis, we trained an LR classifier on the protein-level embeddings to predict protein length. We binned the protein length similar as in Figure 3A and modelled these bins by one-hot encodings.

Indeed, we observed that protein length was modelled by the protein-level embeddings, as reflected by an average ROCAUC score of 0.856 taken over all protein length intervals of the LR classifier (Figure 3C). Specifically, the performance was high for very short or very long proteins, and moderate for proteins with a more average length. Overall, this finding indicated that embeddings still capture relevant information on protein size, even though they are obtained by taking the mean over amino acid embeddings.

### Term-centric performance correlates positively with an increased structural similarity between proteins

Previous research has reported that SeqVec embeddings encode information about secondary and possibly tertiary protein structures, making it likely that proteins with the same structure are close in the embedding space [22, 23]. As an alternative approach to explain the differences in term-centric performance, we hypothesized that differences in the structural similarity between proteins annotated with certain GO terms might be the underlying cause. Specifically, proteins with the same structure tend to perform the same molecular functions and SeqVec embeddings might not contain the useful information to correctly predict GO terms with many annotated structurally dissimilar proteins (yet functionally similar), lowering their term-centric performance [32, 33]. As three-dimensional structures of proteins were not widely available, we retrieved the three next best structural similarity measures from the InterPro database: (i) protein domains (i.e. structurally conserved functional units) (ii) protein families (i.e. evolutionarily related proteins with similar three-dimensional shapes), and (iii) protein superfamilies (i.e. structurally/mechanistically related proteins not necessarily evolutionary related). As even these structural annotations were sparse, we were unable to retrieve annotations for all test proteins.

To quantify the structural similarity between proteins annotated with a certain GO term, we evaluated what percentage of them shared a structural domain, family or superfamily annotation. Note that these annotations are not independent quantities (Spearman correlations; domains-families: 0.68, p-value: 1.6e-29; domains-superfamilies: 0.75, p-value: 5.2e-42; families-superfamilies: 0.49, p-value: 3.3e-14). We observed moderate non-linear positive correlations between the percentage of proteins sharing a domain, family or superfamily and term-centric performance (Spearman correlations: 0.43, p-value: 9.5e-14; 0.37, p-value: 7.4e-13; 0.30, p-value: 5.5e-7, respectively) (Figure 4A). This indicates that indeed the term-centric performance for GO terms with many annotated structurally similar proteins is generally better, which is additional evidence that SeqVec embeddings capture information about protein structures.

**Fig 4.**
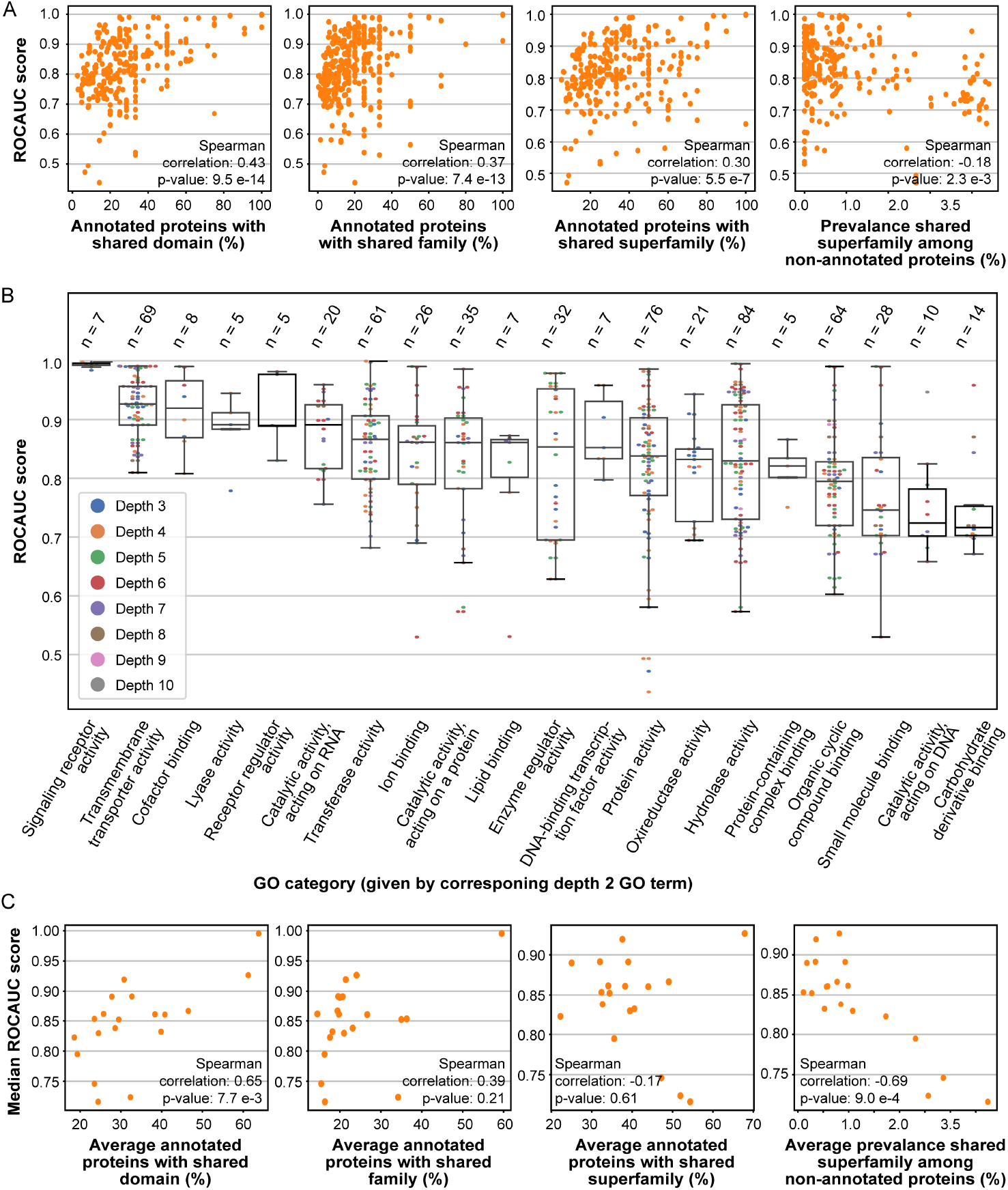
(A) ROCAUC per GO term of the SeqVec-based LR classifier in relation to the structural similarity of proteins annotated with that term. From left to right: the performance in relation to the percentage of annotated proteins with a shared domain, family or superfamily. On the far right: performance in relation to the prevalence of the shared superfamily among the remaining non-annotated proteins. (B) ROCAUC per GO term of the same classifier in relation to GO category. Child GO terms in each GO category are shown as dots with colour indicating their depth. The number of child terms per category is presented at the top. The box-whiskers plots show the standard IQR, median and 1.5*IQR. (C) As in A, but terms are grouped per GO category and the median ROCAUC is shown for each category.

Next, we hypothesised that SeqVec’s ability to model structural similarities might be compromised when a protein is structurally annotated with multiple domains, families or superfamilies as more information needs to be captured in the same 1024-dimensional embedding. However, we observed no statistically significant correlations between the average number of domains, families or superfamilies per annotated protein and term-centric performance, indicating that SeqVec embeddings efficiently modelled multiple functionalities of proteins (figure S.2A).

Finally, as we still observed significant differences in term-centric performance between GO terms with a low percentage of proteins sharing a domain, family or superfamily, we hypothesised that a higher prevalence of the shared domain, family or superfamily among the remaining test proteins could lower performance as they might be predicted as false positives. Indeed, we observed a weak non-linear negative correlation between the prevalence of the shared superfamily for a certain GO term among the remaining test proteins and the term-centric performance (Spearman correlation: −0.18, p-value: 2.3e-3) (Figure 4A). We did not observe a statistically significant correlation between the prevalence of the most shared domain or family and the term-centric performance, possibly due to the generally low prevalence of the most shared domain or family in the remaining population (figure S.2B).

Overall, these correlations hint that some molecular functions are being executed by a wider spectrum of protein structures, thereby lowering the term-centric performance of SeqVec-based molecular function prediction.

### High term-centric performance related to specific types of molecular functions with high structural similarity

To identify the molecular functions with many structurally similar annotated proteins, we created so-called ‘GO categories’. First, we selected all GO terms in the SwissProt dataset with depth two. Next, from this selection, we selected terms with at least five child terms. This resulted in twenty ‘GO categories’ indicated by their depth 2 GO term, thereby indicative of certain types of molecular functions.

We ordered the GO categories based on their median term-centric performance and observed large differences in their term-centric median performance and the spread in performance (Figure 4B). Notably outstanding was the GO category of ‘signalling receptor activity’ with a median performance of ~0.98 and almost no spread. To confirm if indeed the molecular functions of the best performing GO categories were executed by structurally similar proteins, we related the performance of each GO category to the four significant correlations on protein structure similarity measures mentioned in the previous section (see Table S.1). As expected, we observed a strong positive correlation between the average percentage of shared domains among the annotated proteins and the median term-centric performance of the GO category (Spearman correlation: 0.65, p-value: 7.7e-3) (Figure 4C). Additionally, we observed a strong negative correlation between the average prevalence of the shared superfamily among the remaining proteins and the median term-centric performance of the GO category, indicating that proteins with more common superfamilies can generally be predicted worse (Spearman correlation: −0.69, p-value: 9.0e-4). We did not observe a statistically significant correlation between the average percentage of shared families/superfamilies among the annotated proteins and the median term-centric performance of the GO category.

These results suggest that differences in term-centric performance for SeqVec-based molecular function prediction models mainly stem from differences in the structural divergence between proteins executing the same molecular functions.

## Results cross-species function prediction

### Model species selection

We assessed if knowledge about protein function learned in one training species could be generalized to other species. To this end, we trained a SeqVec-based molecular function prediction model on the data of one training species and assessed its performance on the data of several test species with varying evolutionary distance. As proof of principle, we considered seven well-annotated species from different evolutionary classes, phyla and even kingdoms: *Mus musculus* (Mouse), *Rattus norvegicus* (Rat), *Homo sapiens* (Human), *Danio rerio* (Zebrafish), *Caenorhabditis elegans* (C. elegans), *Saccharomyces cerevisiae* (Yeast) and *Arabidopsis thaliana* (A. thaliana) (Table 1A) [34]. As these species have different genome sizes, they have a different number of testable protein sequences. To quantify how well these proteins represented all the molecular functions in the species we calculated the coverage of the gene count, i.e. how many of the protein-coding genes were represented by the proteins (Table 1A). We selected Mouse as the training species, creating an ‘evolutionary staircase’ in which the remaining species had increasing divergence time from Mouse (Figure 5A). To optimally tune and assess a classifier, we split the Mouse data into 8.977 mouse training, 1.801 mouse validation and 1.790 mouse test proteins.

**Fig 5.**
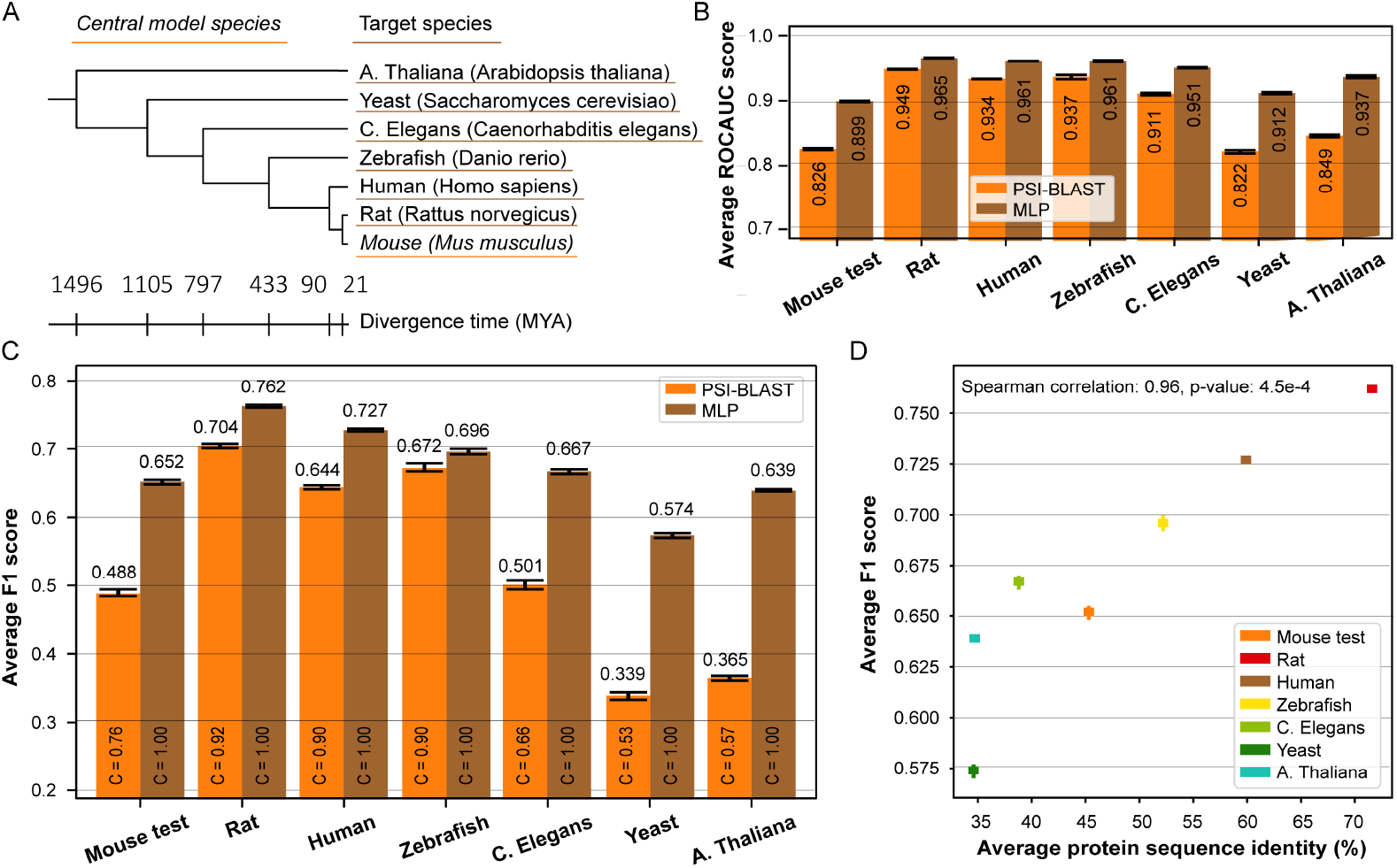
(A) Phylogenetic tree showing evolutionary relation and divergence time between the training species Mouse and the other test species. Tree produced via the PhyloT tool for phylogenetic tree visualisation and divergence times retrieved using the TimeTree tool [36, 37]. (B) Average term-centric ROCAUC over all the GO terms and (C) average protein-centric F1 over all the proteins per species for the MLP classifier (brown). The MLP was trained to predict GO terms. Performance is compared to baseline PSI-BLAST (orange). In (C) the coverage C is shown inside the bars. (D) Average protein-centric performance (F1) over all the proteins per species of the MLP in relation to the average protein sequence identity to the Mouse training set. Errorbars denote 95% confidence intervals estimated using 100 bootstraps.

**Table 1.** List of species used in this study. (A) Taxonomic rankings, number of proteins and gene count coverage for each species. Gene count is a measure for the number of protein-coding genes in the genome [35]. (B) The number of GO terms (evaluated) in each species. During protein-centric evaluation GO terms overlapping with the Mouse training set are considered. During term-centric evaluation only GO terms with at least 3 annotated proteins and overlapping with the Mouse training set are considered. The mouse dataset consists of the mouse training, validation and test sets combined.

| Species | Kingdom | Phylum | Class | # selected proteins | Coverage of gene counts (%) |
| --- | --- | --- | --- | --- | --- |
| Mouse | Animalia | Chordata | Mammalia | 12.568 | 57% |
| Rat | Animalia | Chordata | Mammalia | 6.184 | 29% |
| Human | Animalia | Chordata | Mammalia | 14.823 | 72% |
| Zebrafish | Animalia | Chordata | Actinopterygii | 1.883 | 7% |
| C. elegans | Animalia | Nematoda | Chromadorea | 2.732 | 14% |
| Yeast | Fungi | - | - | 4.358 | 72% |
| A. thaliana | Plantae | - | - | 10.888 | 40% |

**Table 1.** List of species used in this study.
| Species | # GO terms | # GO terms evaluated protein-centric (% of terms) | # GO terms evaluated term-centric (% of terms) |
| --- | --- | --- | --- |
| Mouse | 4.572 | - | - |
| Mouse training | 4.086 | - | - |
| Mouse validation | 2.215 | 1.964 (89%) | 834 (38%) |
| Mouse test | 2.238 | 1.966 (88%) | 855 (38%) |
| Rat | 3.394 | 3.556 (90%) | 1.469 (37%) |
| Human | 4.684 | 4.008 (86%) | 1.523 (33%) |
| Zebrafish | 1.678 | 1.582 (94%) | 654 (39%) |
| C. elegans | 1.968 | 1.816 (92%) | 788 (40%) |
| Yeast | 2.547 | 1.955 (77%) | 840 (33%) |
| A. thaliana | 2.976 | 1.977 (66%) | 899 (30%) |

Besides the different number of test proteins for each species, we note additional differences in the number of GO terms present among the proteins of each species (Table 1B). In practice, one would want to predict as many molecular functions as possible, but for this feasibility study we note two major limitations: (i) we could not test target species GO terms when they were not present among the Mouse training set GO terms (e.g. GO terms related to photosynthesis), and (ii) testing all the Mouse training GO terms in the test species could have predicted annotations for some proteins which we would not be able to reliably validate as data on those functions was lacking. Hence, for protein-centric evaluation (F1-score) we evaluated only the GO terms overlapping between the Mouse training set and the datasets of the test species. For term-centric evaluation (ROCAUC score) we evaluated only GO terms overlapping with the mouse training set and with at least three annotated test proteins (Table 1B).

After the GO term selection process we checked if the selected GO terms for every species were of similar depth as we previously showed a positive correlation between term-centric performance and GO term depth. We observed similar distributions of GO term depth per species, although for at least one species the distribution was significantly different (Chi-square test p-value: 2.7e-23) (Figure S.3A). This was not the case for the depth distributions of GO terms selected for term-centric evaluation (Chi-square test p-value: 0.20) (Figure S.3B). The observed difference in depth distribution for protein-centric evaluation may have a minor influence on differences in performance between test species.

### Protein functions learned in training species are effectively predicted cross-species

We trained an MLP classifier on the embeddings of the Mouse training set. We specifically select the MLP over LR because our interest now lies in best performance. To evaluate protein-centric performance (F1-score), GO term posterior probabilities were converted into predicted binary class labels using a threshold. To mimic a real-case scenario in which no information on the test species is present, we determined this threshold on the mouse validation set and applied it to posterior probabilities for every test species (Figure S.4).

We observed that the MLP outperformed the baseline PSI-BLAST method in all species for both term-centric and protein-centric evaluation (Figure 5B, C). The absolute performance of the MLP decreased with increasing divergence time, yet the decrease was not as severe as for the PSI-BLAST method, effectively increasing the difference by which MLP outperformed PSI-BLAST. We observed deviant behaviour in Mouse, Yeast and A. thaliana as their MLP performance did not follow the trend of ‘increased divergence time, decreased performance’. As the behaviour of Mouse might be caused by splitting the data into a train, validation and test set, we recreated our MLP experiments with Human as the training species as a control experiment (Figure S.5A). We observed a similar trend as before but this time the performance in Human, Yeast and A. thaliana was deviant, indicating that the splitting of the training species into a train, validation and test set was responsible (Figure S.5B, C).

Overall, the results reveal the ability of SeqVec-based molecular function prediction to extract information from one well-annotated training species for predictions in various other eukaryotic species. Additionally, the deviant behaviour of Yeast and A. thaliana indicated that after the divergence time reached some threshold, cross-species SeqVec-based molecular function prediction no longer followed the observed ‘increased divergence time, decreased performance’ trend.

### Cross-species protein-centric performance positively correlates with protein sequence identity

We hypothesized that both the observed decrease in SeqVec-based molecular function performance with increasing divergence time and the deviant performance after dataset splitting could be the result of differences in protein sequence identity to the proteins in the training set. Specifically, the probability of conserved protein function increases with increasing sequence conservation, which could be improving the performance [17].

Indeed, using the PSI-BLAST top hit of every protein to find its sequence identity to the training set, we observed a very strong positive correlation between protein-centric SeqVec-based molecular function prediction performance and average sequence identity per species (Spearman correlation: 0.96, p-value: 4.5e-05) (Figure 5D). From the distributions of the sequence identity, we observed that with increasing divergence time the distributions skewed from high sequence identity to low sequence identity (Figure S.6). The deviant performance of Yeast could partially be explained by the observed correlation, as Yeast had the lowest (but very similar to A. thaliana) average sequence identity. The performance of A. thaliana in relation to its average sequence identity was more corresponding with the other test species, indicating that Yeast is likely breaking the trend.

On the other hand, proteins in the ‘twilight zone’ have below 30% sequence identity resulting in limited structural similarity, decreasing the likelihood of conserved protein function [38, 39]. As the amount of sequence identity and the divergence time between species are related, we repeated the MLP experiments while evaluating only proteins from the twilight zone. While we observed a very similar average sequence identity in all species after the sequence identity constraint, the spread in performance remained large (Figure S.5D). Additionally, the species with the highest sequence identity did no longer have the best performance (Figure S.5E). We no longer observed a statistically significant correlation between performance and sequence identity (Spearman correlation: −0.21, p-value: 0.64). These findings illustrate that for proteins in the twilight zone their sequence identity is no longer indicative of performance. As we observed that Yeast had the highest fraction of proteins in the twilight zone, this possibly explains the low Yeast performance (Figure S.6).

Overall, these observations indicate that divergence time is not in itself a good predictor of accuracy of cross-species function prediction.

### Not-evaluated species-specific GO terms contribute only a few annotations

As cross-species molecular function prediction inevitably limits the number of GO terms that can be predicted in test species, we assessed to what extent this affects the integrity of SeqVec-based molecular function prediction. Specifically, a protein has a certain number of real annotations which are all the annotations present in the species datasets, including the non-evaluated GO terms. Given that we only evaluated overlapping GO terms between training and test species, we calculated the percentage of these real annotations that we were able to predict as a quantity for missed predictions.

In all species, we observed a wide distribution of the predicted percentage of real annotations per protein (Figure S.7A). The distributions were heavily tailed to the high percentages for all test species except the Mouse and Yeast, which previously also showed lower performances. Interestingly, with increasing divergence time, the average percentage of predicted real annotations remained fairly constant, whereas the percentage of real GO terms evaluated decreased with increasing divergence time (Table 1). We hypothesised this observation might be indicating that the filtered-out GO terms were rare terms with few annotated proteins, and hence excluding them from evaluation had an only minor influence on the percentage of true predicted GO terms.

To test this, we calculated the Resnik information content (IC) [40] of each GO term per species. A high IC indicated a rare term in a GO corpus, and a low IC a common term. Indeed, the average IC value of not-evaluated GO terms was high for all species, indicating that the not-evaluated GO terms represent rare molecular functions among the proteins of the test species (Table S.2).

### SeqVec-based function prediction covers much more proteins than established PSI-BLAST method

Besides the quality of molecular function prediction method, its coverage (i.e. the fraction of protein for which at least one prediction was made) is a second most important characteristic.

We observed that the coverage of the classifiers trained using SeqVec embeddings was 1.0 in all experiments (Figure 5B, Figure S.5C). 50-80% of the proteins were assigned a term of depth at least 4 and these percentages decreased for increasing term depth (Figure S.7B). The decrease was faster for the species with worse protein-centric performance. For the PSI-BLAST method, we observed a generally low and decreasing coverage with increasing divergence time, indicating the inability of PSI-BLAST to predict even one molecular function for a large portion of evaluated proteins (Figure 5C). This reduction in coverage likely originated in the decreasing average sequence similarity between the training species and the test species.

### SeqVec-based molecular function prediction reveals specific types of molecular functions executed by structurally conserved proteins across different species

From the characterisation of SeqVec-based molecular function prediction, we know that performance positively correlates with an increased structural similarity between proteins. We figured that we can exploit this property to identify types of molecular functions executed by structurally more conserved proteins. Specifically, if performance remains constant with increasing divergence time, it would indicate that the proteins are structurally conserved. Hence, we compared the performance of each GO category between all evaluated species.

We observed that indeed some molecular functions indicated by their GO category could be predicted with constant performance across all species, such as ‘transmembrane transporter activity’ (Figure 6). Additionally, we observed GO categories with constant performance among the mammal species and decreased performance in the other species such as ‘hydrolase activity’ and ‘catalytic activity, acting on a protein’, indicating that these categories of functions are more structurally conserved among mammals.

**Fig 6.**
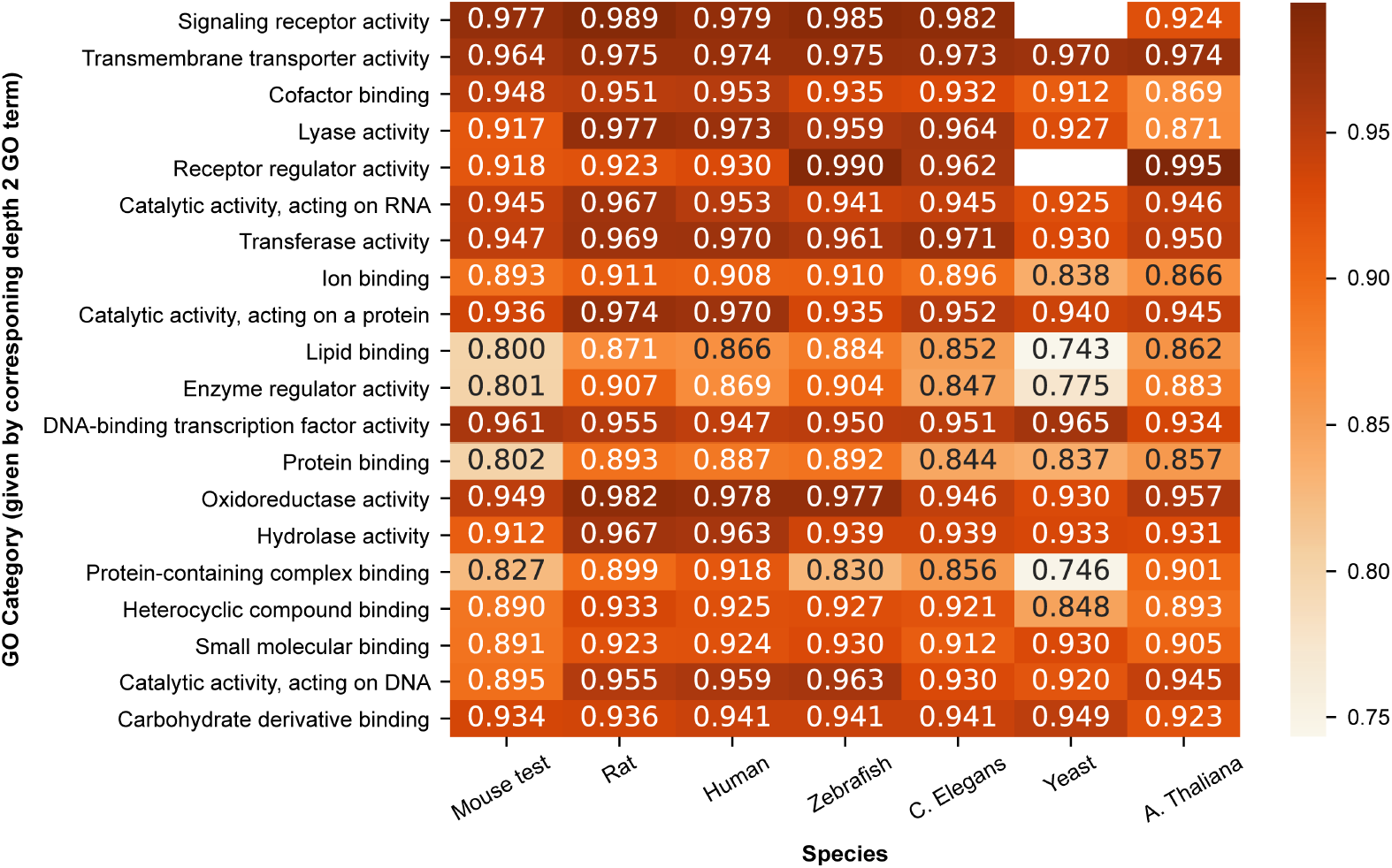
Median term-centric performance (ROCAUC) per GO category per species of the MLP classifier trained using SeqVec protein-level embeddings on the Mouse training dataset. A missing number indicates that a certain GO category was not present among the evaluated proteins.

As we ordered the GO categories the same as in Figure 4 (best to worst performance), it was interesting to observe that the GO categories did not perform similar in the cross-species experiments (i.e. they do not have the same order of performance in the different species). Note, the order was previously determined using the SwissProt dataset (which has maximum 30% pairwise sequence identity), so this dataset disregarded many proteins of certain GO categories with high sequence similarity. Given the discrepancies in order and no restrictions on sequence similarity for the cross-species experiments, these results indicate that in reality a large number of structurally similar proteins likely exist for these categories. Overall, these results reveal a possible application for SeqVec-based molecular function prediction in which conserved protein functions could be identified.

### Protein functions from the GO categories Biological Process and Cellular Component can also effectively be predicted cross-species

To further assess the potential of cross-species SeqVec based protein function prediction, we tried to predict GO terms from the GO categories Biological Process (BP) and Cellular Component (CC) using the same methodology as before. Again using Mouse as the training species, we created BP and CC datasets with protein annotations for the test species (the number of selected proteins and GO terms can be found in Table S.3).

In general, we observed the same trend as before as the absolute performance of the MLP decreased with increasing divergence time (Figure 7, S.8). Again, the decrease was not as severe as for the PSI-BLAST method, effectively increasing the difference by which MLP outperformed PSI-BLAST. Additionally, the coverage of the MLP remains perfect whereas the coverage of PSI-BLAST also declines with increasing divergence time. Outstanding is the protein-centric performance of PSI-BLAST for biological process function prediction (Figure 7A). Here, in species evolutionary close to the training species (i.e. all chordate species) PSI-BLAST outperforms the MLP although its performance declines rapidly with increasing divergence time. For the term-centric performance, however, we observed that the MLP outperformed the baseline PSI-BLAST method in almost all species for both the biological process and cellular component predictions (Figure S.8A, B). This is in line with what was previously observed for molecular function predictions.

**Fig 7.**
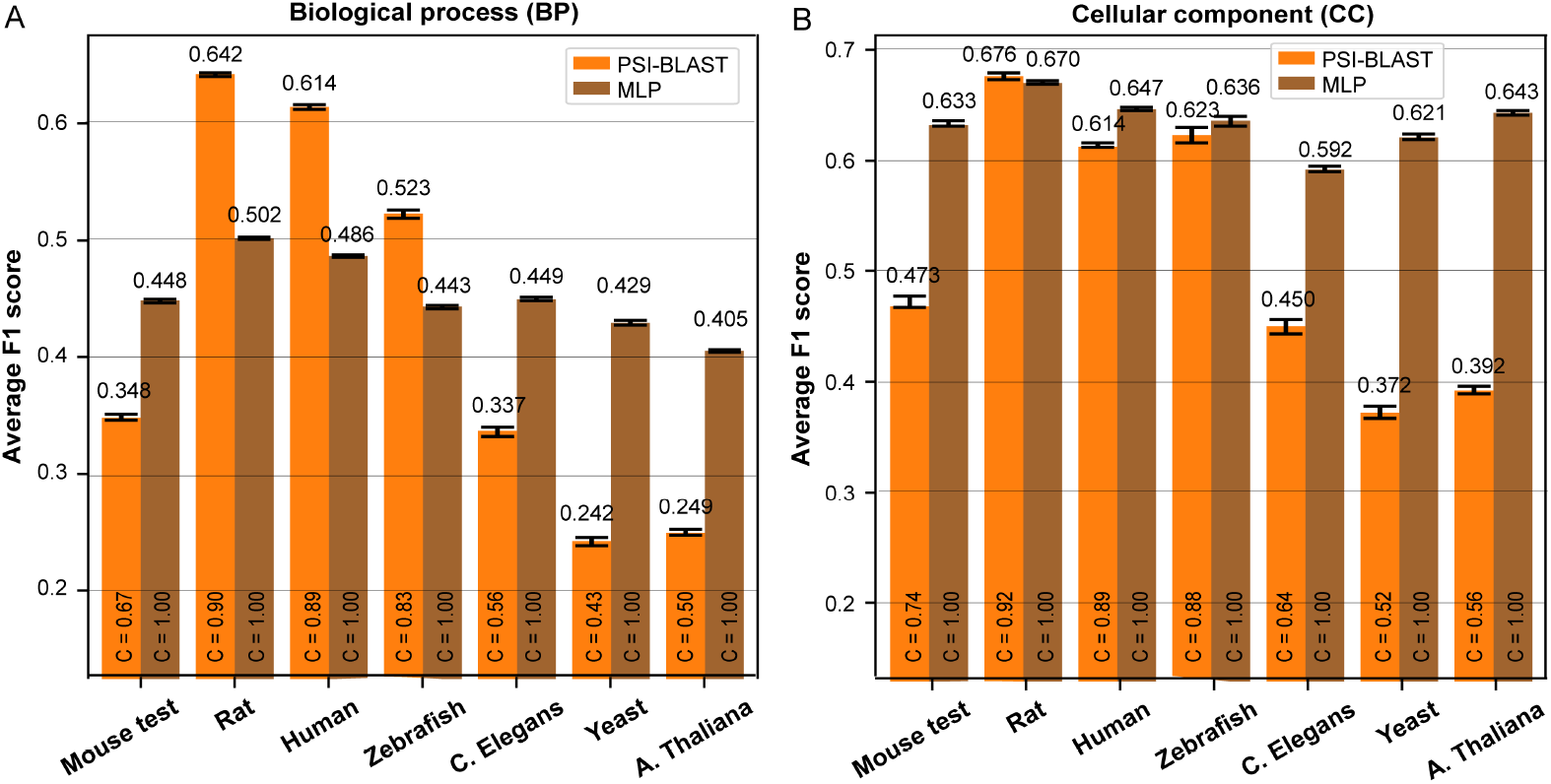
The average protein-centric F1 performance over all the proteins per species for the MLP classifier (brown) for (A) biological process GO terms and (B) cellular component GO terms. Performance is compared to baseline PSI-BLAST (orange). The coverage C is shown inside the bars. Errorbars denote 95% confidence intervals estimated using 100 bootstraps.

Although no further experiments were done on the prediction of biological process and cellular component GO terms, these results further support the potential for SeqVec-based protein function prediction in practical applications.

## Discussion

### Performance of SeqVec-based molecular function prediction models dominated by the level of structurally conserved proteins

Protein-level embeddings from the SeqVec model are effective tools in the task of molecular function prediction, reaching competitive performance to many state-of-the-art sequence-based molecular function prediction methods [22]. Here, we shed light on the ‘black box’ of SeqVec performance and built upon previous work by characterising the performance of a simple LR classifier trained using SeqVec embeddings. We observed a (i) weak positive correlation between term-centric performance and GO term depth, (ii) weak negative correlation between term-centric performance and the number of training proteins per GO term, (iii) weak positive correlation between protein-centric performance and protein length, and (iv) moderate positive correlation between term-centric performance and structural similarity between annotated proteins.

Fundamental to the GO hierarchy, GO terms close to the root annotate a wide variety of proteins compared to more specific divergent functions lower in the GO hierarchy. The first and second observation suggest that inherent to this GO hierarchy, GO terms closer to the root can generally be predicted with lower term-centric performance than GO terms far from the root. This implies that the performance of SeqVec-based molecular function prediction is sensitive to a training set containing proteins roughly similar on a large functional scale, yet still distinct on a smaller scale. A possible countermeasure is offered by ‘projected predictions’ to correct predicted probabilities to respect the GO hierarchy [15]. Specifically, the probability of a protein being annotated with a term close to the root (e.g. ‘binding’) should never be lower than the probability of being annotated with a child term of it (e.g. ‘DNA binding’). In theory, this could improve the predictions of GO terms close to the root, although it will be less effective for GO terms with many false positives which likely already have high predicted probability scores.

The third observation, a positive correlation between protein-centric performance and protein length, was against our expectation of longer proteins having a worse protein-centric performance. To explain the observed behaviour, we suspected that the length of the protein is somehow encoded in SeqVec embeddings. Specifically, as long proteins were relatively scarce in our dataset, they could be easier to predict by having similar embeddings to other long proteins with similar function in the training set. Indeed, we showed that protein length can effectively be predicted from SeqVec embeddings, thereby presenting itself as a protein characteristic that counters the expected decrease in performance for longer proteins. Interestingly, this relevant information on protein length is present in protein-level embeddings, even though they are obtained by taking the mean over amino acid embeddings. This is in line with results from NLP, where sentence embeddings have been shown to be predictive of sentence length, even by averaging the embeddings of all words in a sentence [41].

As we observed large differences in performance between GO terms with the same depth, we hypothesised that some molecular functions are somehow easier to predict than others. We investigated the possible influence of differences in the structural similarity between annotated proteins on term-centric performance as we hypothesised that GO terms with structurally diverse proteins might be harder to predict. Specifically, if SeqVec embeddings can indeed accurately model protein structures, structurally similar proteins might have similar embeddings and hence a higher probability of being assigned the same molecular function [22, 23]. Indeed, the fourth observation confirmed that the term-centric performance for GO terms with many annotated structurally similar proteins is generally better. Additionally, we showed that having multiple protein domains per protein does not interfere with SeqVec’s ability to model structural similarities between proteins. Hence, the SeqVec model is capable of modelling multiple functionalities of proteins in one embedding. Given that SeqVec is trained using biLSTM layers, this observation hints that SeqVec might be able to recognize protein domains in protein sequences which could be the underlying mechanism by which it is capable of modelling multiple functionalities of proteins. Moreover, we showed that having a higher prevalence of the shared superfamily among the remaining protein population lowered term-centric performance. The latter observation is in line with our previous notion that SeqVec-based molecular function prediction performance suffers from having to predict proteins with a similar function at the broad scale yet a distinct specific function. Overall, these observations hinted that some molecular functions are executed by a wider spectrum of protein structures, thereby decreasing the predictive power of SeqVec-based molecular function prediction. Of note, we were unable to consider all test proteins in the experiments on structural similarity as structural annotations were sparse. Nevertheless, we do not expect large differences in the observed findings if all structural annotations would be available.

Using these insights, we identified specific groups of molecular functions executed by more structurally similar proteins. As expected, this higher amount of structural similarity was positively correlated with the median term-centric performance of the GO category, i.e. the group of GO terms with some broad similar molecular function. The higher structural similarity in some GO categories might be explained from an evolutionary perspective: the proteins that execute the molecular functions of well-performing GO categories seem to be more structurally conserved. For instance, our best performing GO category was ‘signalling receptor activity’, and the Par proteins, GTPases, kinases, and phosphoinositides that participate in signalling pathways are highly conserved over diverse species [42]. On the other hand, our worst-performing GO category, ‘carbohydrate derivative binding’, is executed by proteins with a high degree of complexity and heterogeneity, as reflected by the fact that the proteins are grouped into 45 protein families [43, 44]. Therefore, the performance of SeqVec-based molecular function prediction might be indicative of groups of conserved proteins. For instance, we observed that even within the median performing GO categories some GO terms do have high performance, and these more specific molecular functions might be executed by a group of conserved proteins.

Considering all the observations on ‘increased structural similarity, increased performance’ of SeqVec-based molecular function prediction, the SeqVec embeddings of structurally similar proteins are likely also similar. Specifically, these embeddings are likely somehow more similar in the embeddings space, and hence more likely to receive the same functional annotation, increasing term-centric performance. Thereby, we support the claims that SeqVec embeddings somehow must model secondary structure and likely also tertiary structure of the proteins [22, 23].

Overall, these results indicate that differences in term-centric performance for SeqVec-based molecular function prediction models can be partly explained by differences in structural divergence between proteins executing the same molecular functions, and to a lesser extent by GO term characteristics. If this finding can be replicated in other sequence-based function predictors, it might hint that structural divergence should be taken into account when evaluating predicted annotations. This could augment measures of information content, rewarding accurate predictions on terms that are harder to infer because of the large diversity of proteins assigned to them given the number of proteins these term annotate.

### Cross-species SeqVec-based molecular function prediction is possible and offers many fruitful applications in practice

Our work provides a novel evaluation scheme to molecular function prediction based on the annotated protein sequence data of merely one training species. Using the methodology of SeqVec-based molecular function prediction in a transfer learning task, the model effectively extracted information on protein functions from one training species to make predictions in various other eukaryotic species. This ability to generalize learned protein functions across different kingdoms shows that the trends found by the neural network (both the SeqVec model and the MLP classifier) not only hold for the proteins of the training species but are conserved throughout the eukaryotic domain of life. This confirms our hypothesis that SeqVec-based molecular function prediction is to some extent independent to the context of species and substantiate SeqVec’s capability to model underlying protein principles (i.e. the ‘syntax’ of proteins) [23].

We showed that the absolute performance of SeqVec-based molecular function prediction decreased with increasing divergence time, although it was not as severe as for the baseline PSI-BLAST method, effectively increasing the difference by which SeqVec-based molecular function prediction outperformed PSI-BLAST. We correlated this decrease in performance to a decrease in average sequence identity between the training species and the test species. One explanation to this observation comes from the rule of thumb ‘increased sequence identity, increased likelihood structural similarity and hence increased likelihood functional similarity’ between proteins [17, 32]. We previously noted that SeqVec embeddings of (structurally) similar proteins are likely similar in embeddings space, and hence more likely to receive the same molecular function, explaining the observed behaviour. However, if functionally similar proteins are indeed similar in embedding space, one might not expect a substantial decrease in performance with increasing divergence time. Instead of the entire embeddings of (structurally) similar proteins being similar in embeddings space, it could be that certain embedding features correlate with specific molecular functions. These specific embedding dimensions would arguably be more similar between proteins with a higher sequence identity, thereby explaining the observed decrease in performance with increasing divergence time.

However, we note that for proteins in the twilight zone, i.e. with maximum 30% sequence identity, the correlation between performance and sequence identity disappeared. We attributed this to the fact that the relationship between sequence identity and structural similarity vanishes in the twilight zone, potentially lowering the performance if proteins become structurally divergent [39]. With Yeast and A. thaliana both having a significant proportion of their proteins with a lower than 30% sequence identity, the performance of cross-species function prediction becomes more dependent on the randomness of evolution, i.e. how many proteins will be structurally divergent by evolution, thereby lowering the performance of SeqVec-based molecular function prediction. This is illustrated by the performance of Yeast, the species with the highest fraction of proteins in the twilight zone and the worst performance, while not being the furthest in divergence time. Overall, this indicates that after some threshold in divergence time from the training species, the molecular function prediction will disobey the observed ‘increased divergence time, decreased performance’ trend. Hence, for high-performance in the test species, the training species should be reasonably evolutionary close. Ideally, one model species per kingdom (Plantae, Fungi, Bacteria) could deal with this problem.

Therefore, a fruitful application of our approach arises in cases where limited annotated protein training data is available per taxonomic kingdom. Using the direct transfer approach, we can train a classifier on one training species and directly apply it to protein sequence data of test species. For instance, the UniProtKB/SwissProt knowledgebase has started the Plant Proteome Annotation Program (PPAP) in 2009 to get more annotations on two model plant species, Arabidopsis thaliana and Oryza sativa, which could function as the training species in the plant kingdom [45].

Additionally, we showed the possible application to identify structurally conserved proteins across different species. We previously concluded that the performance of SeqVec-based molecular function is indicative of the amount of structural similarity between the proteins annotated with a certain molecular function, making a constant prediction performance across species indicative of conserved proteins. This makes cross-species SeqVec-based molecular function prediction promising to help formulate biological hypotheses and guide wet-lab experiments. Particularly, we highlight its potential as (i) the coverage of the method is perfect, predicting at least one molecular function for every protein, (ii) for a large fraction of the proteins, this prediction will be of at least depth 4, thereby directly providing significant insight into the molecular function of the protein with unknown function, and (iii) the non-evaluated GO terms between species represent rare molecular functions among the proteins, thereby not contributing much protein functions to molecular function prediction, indicating that cross-species SeqVec-based molecular function prediction covers a large portion of available molecular functions.

Another application arises with the discoveries of novel protein functions. A recent study on the proteomes of 100 species has identified many highly expressed proteins without any functional annotation or sequence homology to proteins with known annotations [3]. It is proposed that the exploration of this dark proteome could reveal essential functions for the species which could be of biological or biotechnological interest. Some novel experimentally identified functions in one training species could quickly be generalized by our cross-species approach to other species, thereby making cross-species SeqVec-based molecular function prediction an effective tool to transfer functional annotations.

Finally, we also evaluated cross-species SeqVec-based protein function prediction for the GO terms on Cellular Component (CC) and Biological Function (BF), the remaining categories of the Gene Ontology. For CC we observed the same behavioral trends in performance as for molecular function predictions. However, a small drop in performance was observed when the test species became evolutionary distant enough (i.e. were from a different taxonomic phylum) from the training species. This behavior is to be expected as CC refers to the cellular anatomy which can highly differ between species from different kingdoms/phyla. For instance, plant cells have chloroplasts, cell walls and a large vacuole making the significantly different from animal cells. In turn, Yeast cells do have a cell wall, but it is not made from cellulose like the cell walls from plants. Therefore, the extensive prediction of cellular components between species would likely be kingdom specific, highlighting the importance of one model species per taxonomic kingdom. BP GO terms describe describe broader functions, much like cellular functions, which already have been shown to generalize between eukaryotes [28].

We observed as well that this is possible. Nevertheless, the baseline method PSI-BLAST protein-centric outperformed the SeqVec method in evolutionary close species, although at a lower coverage. This outperforming, however, was not observed for term-centric performance. This indicates that BP GO terms with many annotations (i.e. terms close to the root) are likely better predicted using PSI-BLAST by contributing many true annotations to proteins which lowers protein-centric performance. Although we did not investigate this further, this could again indicate that SeqVec-based function prediction is better at predicting more specific GO terms which is arguably a desired property. Overall, SeqVec-based function prediction can be employed in predicting molecular functions, cellular components and biological functions, thereby strengthening its practical applications potential.

Future work on cross-species functional annotation should investigate potential domain shifts. As protein function is determined by the context of species, this context could be projected into the classifier during supervised training on only Mouse proteins [15]. This could make the cross-species SeqVec-based molecular function prediction model sensitive to changes in feature distributions between training (source) and test (target) proteins [46]. The more divergent the training species and the test species are, the bigger the influence of this domain shift would be. To counteract this domain shift, several supervised domain adaptation methods have been proposed for the case when (little) labelled target data is available. For certain species, this would most likely increase the SeqVec-based molecular function prediction performance, although most species would require unsupervised domain adaptation due to the lack of available labelled data. Luckily, there are effective methods available for this. For instance, the CORrelation ALignment (CORAL) method would minimize domain shift by aligning the second-order statistics of the distributions of the Mouse and test species protein-level embeddings [46]. As we implemented no such method, it would be interesting to evaluate how performance changes using this method to adapt to the context of species.

In addition, most molecular function prediction methods train on a labelled dataset containing protein sequences from many different species. We note the substantial animal bias in these datasets: for illustration, almost half the proteins from the SwissProt database with a molecular function annotation originate from mammals. As the context of the species determines protein function, this animal bias propagates during supervised training on a dataset that contains all proteins from SwissProt. An interesting outstanding experiment would be to predict protein functions of a plant species with a function prediction model trained using either the labelled data of one other plant species (our cross-species approach) or on the labelled data of all known proteins (a SwissProt dataset). Comparing the performance might indicate whether or not the animal bias reduces molecular function prediction power in cases like this. And if so, this would substantially invigorate the power of our cross-species approach, by which training on a model species in each kingdom should improve supervised training.

All in all, we present a novel, data-undemanding protein function prediction evaluation scheme that relies on the availability of merely one adequately annotated model species per evolutionary kingdom and uses the methodology of SeqVec-based molecular function prediction. We identify multiple fruitful applications for this approach, thereby proving itself an effective method for molecular function prediction in inadequately annotated species from understudied taxonomic kingdoms.

## Materials & Methods

### Datasets: SwissProt and cross-species

#### SwissProt dataset

To characterise and improve the performance of SeqVec-based molecular function prediction, we used the SwissProt dataset from previous work [22]. In short, this dataset contained labelled protein sequence data from the SwissProt database for a selection of proteins with a sequence length in the range [40, 1000]. Every protein had at least one functional GO term annotation from the Molecular Function Ontology (MFO) obtained by non-computational means, i.e. with one of the following evidence codes: ‘EXP’, ‘IDA’, ‘IPI’, ‘IMP’, ‘IGI’, ‘IEP’, ‘HTP’, ‘HDA’, ‘HMP’, ‘HGI’, ‘HEP’, ‘IBA’, ‘IBD’, ‘IKR’, ‘IRD’, ‘IC’, ‘TAS’. Data on the annotations of 441 GO terms with at least 40 positive examples in the training set and at least 5 positive examples in the validation and test set was available. The dataset contained 63,994 training, 8,004 validation and 3,530 test proteins with at most 95% sequence identity to each other. Proteins in the test set had the additional constrain of at most 30% sequence identity to each other and proteins in the training set.

#### Structural similarity data for SwissProt dataset

To measure structural similarity among proteins in the SwissProt dataset, we retrieved all available domain, family and superfamily annotations from the InterPro database. We were able to obtain at least one domain, family or superfamily annotation for 1631, 2210 or 1687 out of the 3530 test proteins, respectively. To prevent calculating statistics over just one or two structurally annotated proteins per GO term, we considered only GO terms with at least 50% of its functionally annotated proteins also being structurally annotated. As a result, we evaluated 278, 355 or 277 out of the 441 GO terms for the domain, family or superfamily similarity, respectively.

#### Cross-species datasets

We tested the ability to transfer knowledge on molecular function between different species using data from seven model species: *Mus musculus* (Mouse), *Rattus norvegicus* (Rat), *Homo sapiens* (Human), *Danio rerio* (Zebrafish), *Caenorhabditis elegans* (C. elegans), *Saccharomyces cerevisiae* (Yeast) and *Arabidopsis thaliana* (A. thaliana). Independent for each of the model species, we retrieved data on the sequence and MFO of proteins from the Swiss-Prot Database, only including proteins with at least one MFO annotation obtained using non-computational means. We retrieved gene counts from the Uniprot reference proteomes [35]. Since mouse was selected as the training species, the mouse data was split into a train, validation and test set using a stratified multi-label split to preserve as many overlapping GO terms as possible between them [47]. This resulted in 8.977 mouse training, 1.801 mouse validation and 1.790 mouse test proteins (ratio of 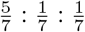 respectively). An overview of the taxonomic classification, the amount of selected proteins and gene coverage per species is provided in Table 1A.

#### Amino acid-level and protein-level embeddings

We represented amino acids in the form of SeqVec embeddings [23]. For every amino acid *n* in the protein sequence, we extracted the *d* = 1024-dimensional embeddings 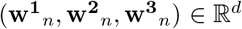 from the three layers of the SeqVec model (Figure 1B). As proposed by [23], we summed these three embeddings component-wise using:

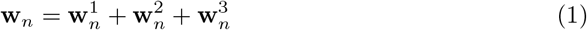

to obtain an amino acid-level embedding 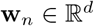.

Using the amino acid-level embeddings, we represented protein sequences with protein-level embeddings [23]. For a protein of length *M*, we calculated the protein-level embedding as the component-wise mean over the sequence of amino acid-level embeddings (**w**_1_,…, **w**_*M*_). Specifically, for every protein we obtained the concise matrix 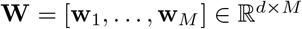 and calculated the vector **v**_1_(**W**) in 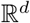 whose *d* components were the component-wise arithmetic mean using:

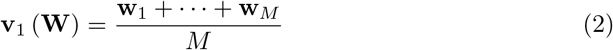

This operation summarized the amino acid sequence of variable length M into a fixed-sized vector **v**_1_(**W**) (Figure 1C). Each of 1,024 protein-level features was standardized to zero mean and unit variance using the training set.

### Molecular function prediction models

#### Models for the SwissProt dataset

We characterised the performance of SeqVec-based molecular function prediction using a Logistic Regression (LR) classifier trained using the protein-level embeddings. The trained classifier predicted for each test protein the probability ∈ [0, 1] of being annotated with a certain GO term.

We trained an independent LR for every GO term using L2 regularization and Stochastic Gradient Descent (SGD) to accelerate the training process. To tune the penalty coefficient λ, we tested the values 10^*x*^ with x ∈ [2,1,0,−1,−2,−3] using the SwissProt validation set. The optimal value was determined jointly over all the 441 GO terms by the highest average ROCAUC score (term-centric evaluation).

#### Model for the cross-species datasets

For the cross-species experiments, we trained an MLP with one hidden layer with 512 nodes followed by a ReLu activation function. We applied a dropout to the hidden layer of 30% to prevent overfitting [48]. The input layer contained a number of nodes equal to the dimension of the input protein-level embeddings, i.e. 1024. The output layer contained 4.086 nodes for all the 4.086 GO terms in the mouse training set, followed by a Sigmoid activation function, ensuring the MLP outputs are in the range [0, 1]. We trained the MLP in a mini-batch mode (size 64) for 100 epochs using the binary cross entropy averaged over all the GO terms as a loss function. We used the Adam optimizer [49] for parameter updating at an initial learning rate of 5 · 10^−4^ that was reduced by a factor of 10 whenever the validation loss did not improve for five consecutive epochs. To obtain the optimally trained MLP model, we selected an independent model for term-centric evaluation and protein-centric evaluation determined by the highest validation performance using the Mouse validation set. Additionally, after the MLP predicted the probabilities of GO term annotations, we propagated them to respect the GO hierarchy. Specifically, each parent term in the GO hierarchy received the highest probability score among its child terms, if and only if this score was higher than its own predicted probability score. This GO hierarchy correction step was not done for the experiments using the SwissProt dataset.

### Protein length prediction model

To access if protein-level embeddings modelled protein length, we trained a LR classifier to predict protein length. We used one-hot encoding to model protein length in bins. The LR was implemented as described above. To tune the penalty coefficient λ, we tested the values 10^*x*^ with x ∈ [2,1,0,−1,−2,−3,−4,−5] using the SwissProt validation set. The optimal value was determined jointly over all the ten protein length intervals by the highest average ROCAUC score (term-centric evaluation).

### Baseline method PSI-BLAST

We used PSI-BLAST with three iterations as a baseline method in the cross-species experiments [16, 50]. We considered all PSI-BLAST hits to the target protein to obtain predicted probability scores for the GO terms as suggested by [15]. In brief, we annotated each protein with all the GO terms present among all the PSI-BLAST hits. The predicted probability given to each annotation was the maximum sequence identity between the target protein and all the PSI-BLAST hit proteins annotated with that term.

### Performance evaluation

We evaluated all models using the protein-centric F1 score and term centric ROCAUC, whose definitions can be found in the supplementary material. We estimated 95% confidence intervals using bootstrapping. W obtained a stratified resampled set from the test set with size equal to the original test set. Subsequently, we calculated the evaluation metrics on the resampled set, repeating this process 100 times using a distinct random state. We executed every bootstrapping process on the different datasets with the same random states to enable comparison between them.

#### GO term selection for cross-species evaluation of models

The cross-species datasets differed in the number of unique GO terms present among the selected proteins. To deal with these differences, we evaluated only GO terms overlapping between the training species (Mouse) and the test species in case of protein-centric evaluation. For term-centric evaluation we had additional criteria as the proper calculation of ROCAUC scores needs enough positive examples for each GO term. To this end, we selected GO terms with at least 5 annotated proteins in the mouse training set and at least 3 annotated proteins in the mouse validation, mouse test and the test species test sets. There were 1530 unique GO terms in the mouse training set with ≥5 annotated proteins. Again, we only evaluated GO terms from the test species overlapping with this selection. An overview of the amount of GO terms per species and evaluation metric is provided in Table 1B.

## Acknowledgements

The authors would like to thank Roeland van Ham and Amelia Villegas-Morcillo for the useful discussions on cross-species prediction and protein embeddings, respectively. We thank Raman van Wee for critical reading and feedback on the manuscript.

## Supporting information

S1 Text. Supplementary Material.

## Author Contributions

Formal Analysis, Software, Writing - Original Draft Preparation: IvB Conceptualization, Methodology, Supervision, Writing - Review & Editing: SM, MR

