## Supplementary material for "The power of universal contextualised protein embeddings in cross-species protein function prediction"

Irene van den Bent, Stavros Makrodimitris and Marcel J.T. Reinders [1]

[1]To whom correspondence should be addressed.

#### Characterization

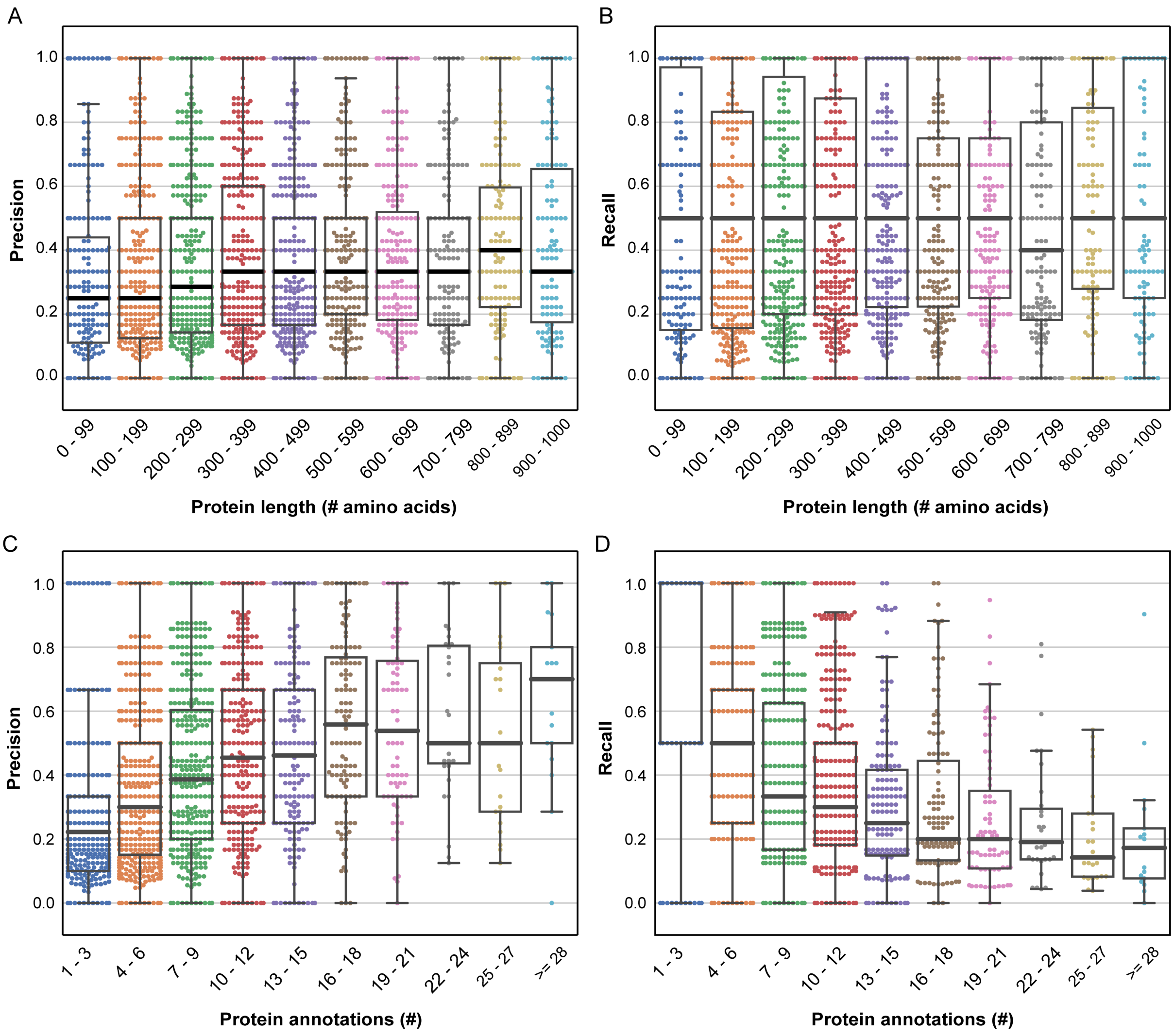


Figure S.1: Protein-centric (A, C) precision and (B, D) recall per protein of LR classifier trained using baseline SeqVec protein-level embeddings on the SwissProt dataset with at most 30% sequence identity in relation to (A, B) protein sequence length and (C, D) the number of protein annotations. The LR was trained to predict GO terms. The box-whiskers plots show the standard IQR, median and whiskers.

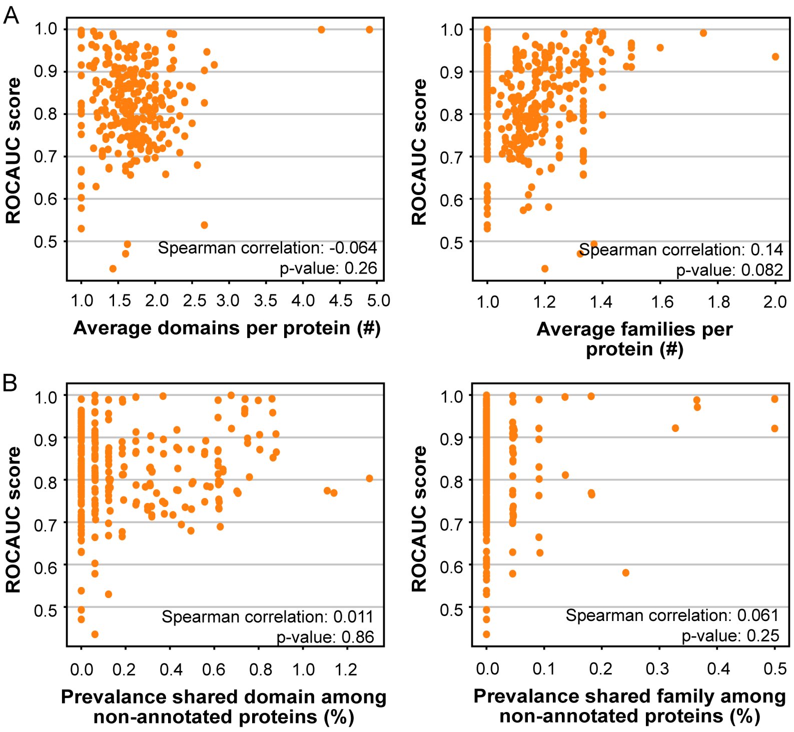


Figure S.2: Term-centric performance (ROCAUC) per GO term of LR classifier trained using baseline SeqVec protein-level embeddings on the SwissProt dataset with at most 30% sequence identity in relation to (A) average number of domain, family or superfamily annotations per protein and (B) prevalence of the shared domain or superfamily among the remaining non-annotated proteins. The LR was trained to predict GO terms. The Spearman correlations and corresponding p-values between term-centric performance and the structural similarity measures are shown. Note that not all GO terms from the SwissProt dataset were evaluated due to a lack of structural annotations (278, 355 or 277 out of the 441 GO terms for the domain, family or superfamily similarity, respectively).

| GO Category | Child GO terms (#) | Median performance (ROCAUC) | Average annotated proteins with shared domain (%) | Average annotated proteins with shared family (%) | Average annotated proteins with shared superfamily (%) | Average prevalence shared superfamily among non-annotated proteins (%) |
|---|---|---|---|---|---|---|
| Signaling receptor activity | 7 | 0.995 (0.000) | 66 | 60 | NaN | NaN |
| Transmembrane transporter activity | 69 | 0.927 (0.003) | 61 | 24 | 68 | 0.81 |
| Cofactor binding | 8 | 0.919 (0.004) | 31 | 21 | 38 | 0.43 |
| Lyase activity | 5 | 0.891 (0.004) | 33 | 20 | 32 | 0.34 |
| Receptor regulator activity | 5 | 0.891 (0.004) | NaN | 20 | 25 | 0.71 |
| Catalytic activity, acting on RNA | 20 | 0.890 (0.004) | 28 | 21 | 39 | 0.93 |
| Transferase activity | 61 | 0.866 (0.006) | 47 | 19 | 49 | 0.77 |
| Ion binding | 26 | 0.861 (0.010) | 26 | 14 | 34 | 1.0 |
| Catalytic activity, acting on a protein | 35 | 0.861 (0.013) | 38 | 20 | 38 | 0.48 |
| Lipid binding | 7 | 0.860 (0.015) | 41 | 27 | 44 | 0.56 |
| Enzyme regulator activity | 32 | 0.853 (0.016) | 24 | 36 | 32 | 0.01 |
| DNA-binding transcription factor activity | 7 | 0.852 (0.004) | 29 | 35 | 34 | 0.27 |
| Protein activity | 76 | 0.838 (0.015) | 29 | 23 | 33 | 0.87 |
| Oxireductase activity | 21 | 0.832 (0.006) | 40 | 18 | 40 | 0.46 |
| Hydrolase activity | 84 | 0.830 (0.012) | 25 | 21 | 39 | 1.0 |
| Protein-containing complex binding | 5 | 0.823 (0.002) | 19 | 18 | 22 | 1.8 |
| Organic cyclic compound binding | 64 | 0.794 (0.008) | 19 | 16 | 36 | 2.4 |
| Small molecule binding | 28 | 0.746 (0.011) | 24 | 15 | 47 | 2.9 |
| Catalytic activity, acting on DNA | 10 | 0.724 (0.007) | 32 | 34 | 52 | 2.6 |
| Carbohydrate derivative binding | 14 | 0.716 (0.007) | 24 | 16 | 54 | 3.5 |

Table S.1: Overview of structural similarity measures per GO category. NaN values indicate a lack of data. Parentheses behind the median term-centric performance (ROCAUC) correspond to the standard deviation of the median performance. The average percentage of annotated proteins with a shared domain, family or superfamily and the average prevalence of the shared superfamily among the non-annotated proteins was calculated over all the child terms in the GO category.

### Cross-species function prediction

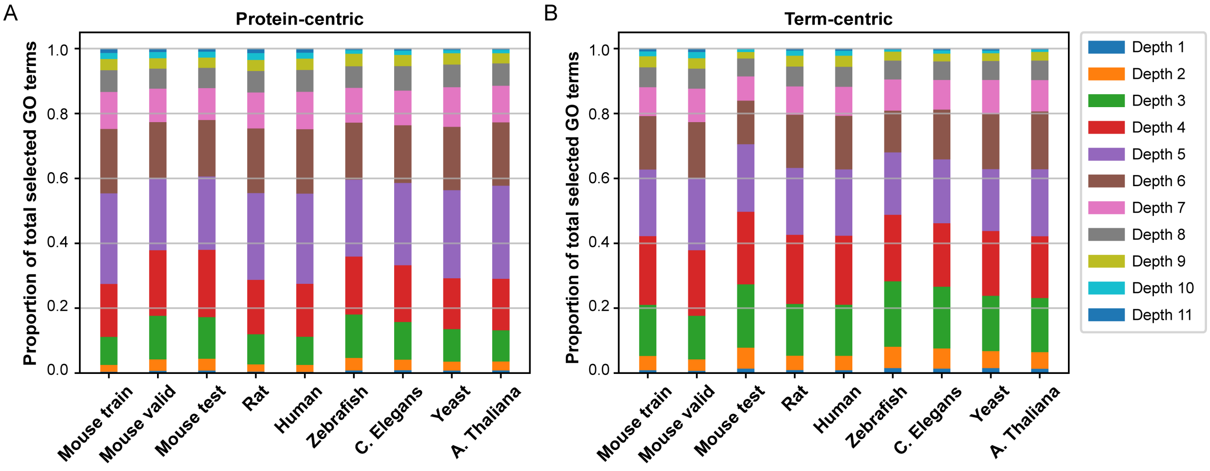


Figure S.3: Distribution of the depth of selected GO terms per species dataset for (A) protein-centric evaluation and (B) term-centric evaluation. Note, in each species a different total number of selected GO terms is present. For at least one species the distribution of GO term depth for protein-centric evaluation was significantly different (Chi-square test p-value: 2.7e-23). This was not the case for the depth distributions of GO terms selected for term-centric evaluation (Chi-square test p-value: 0.20).

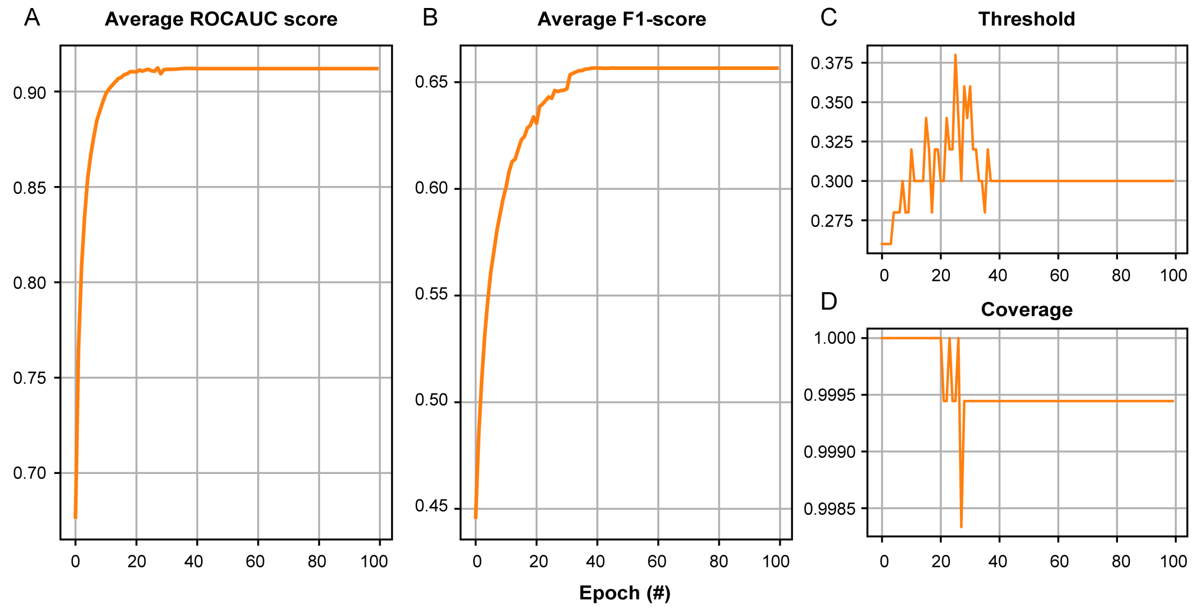


Figure S.4: Average (A) term-centric (ROCAUC) and (B) protein-centric (F1) validation set performance over all the GO terms for MLP classifier trained using baseline embeddings on the Mouse training set. The optimally trained MLP model was selected based on the highest average ROCAUC scores over all the GO terms and the highest F1 score over all the proteins. Note, the MLP was tuned independently for protein-centric and term-centric performance. Each Fmax value was caulated using a certain (C) the threshold on predicted class label probabilities to obtain binary class label predictions. A certain (D) coverage was the result of this threshold. Coverage is the proportion of proteins with at least one predicted GO annotation for the set threshold given the total number of proteins to classify.

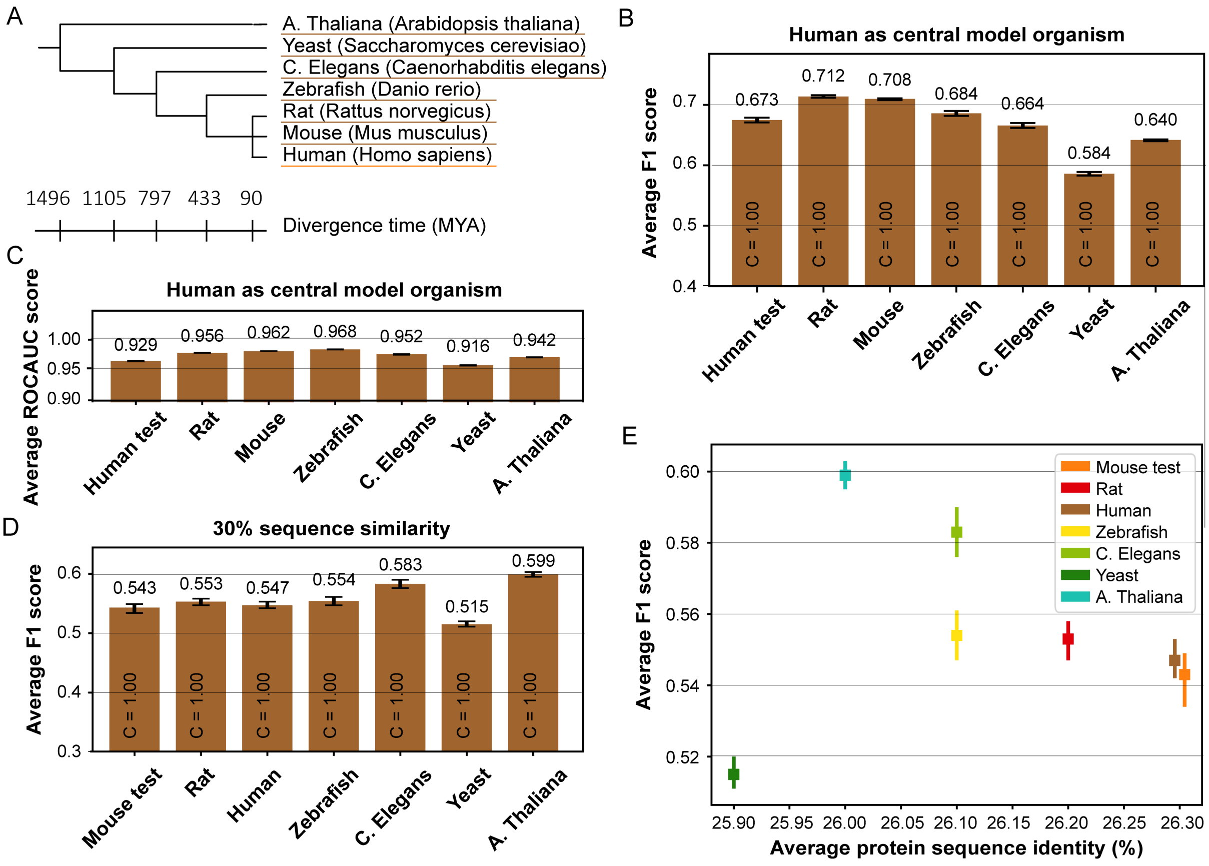


Figure S.5: (A) Phylogenetic tree showing evolutionary relation and divergence time between the train species Human and the other test species, both indicated by colour. Tree produced via the PhyloT tool for phylogenetic tree visualisation and divergence times retrieved using the TimeTree tool. (B) The average protein-centric (F1) performance over all the proteins and (C) average term-centric (ROCAUC) performance over all the GO terms per species of MLP classifier trained using baseline SeqVec protein-level embeddings on the Human training dataset. The MLP was trained to predict GO terms. (D) The average protein-centric (F1) performance over proteins with a maximum 30% sequence identity to the Mouse training set per species of MLP classifier trained using baseline SeqVec protein-level embeddings on the Mouse training dataset. The MLP was trained to predict GO terms. In (B, D) the coverage is shown inside the bars. (E) Average protein-centric performance (F1) over proteins with a maximum 30% sequence identity to the Mouse training set per species of the same MLP classifier in relation to the average protein sequence identity to the Mouse training set. Errorbars denote 95% confidence intervals.

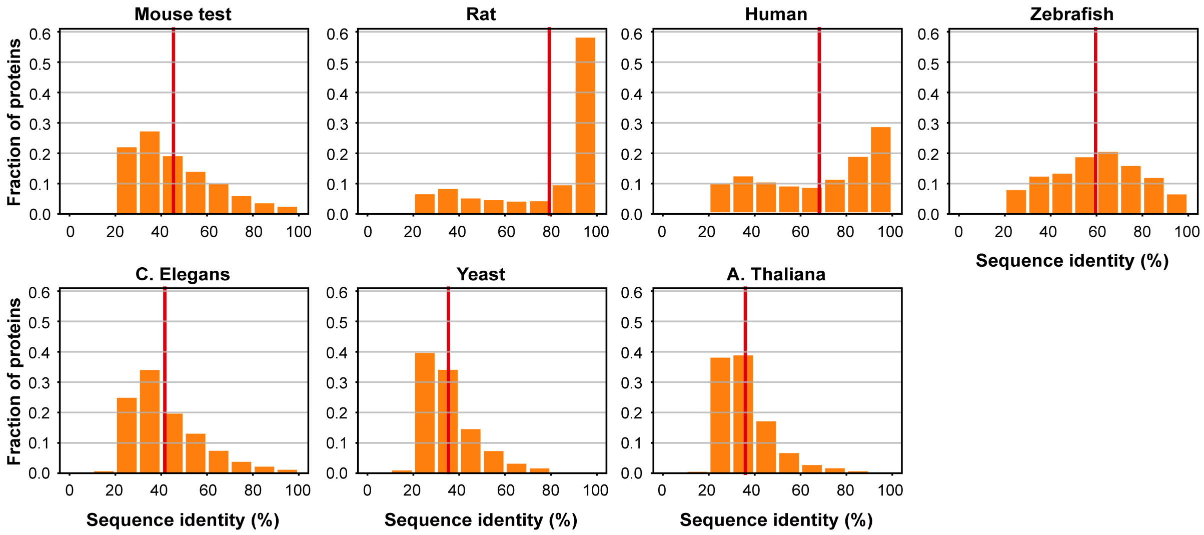


Figure S.6: Distributions of the percentage of sequence identity to the Mouse training set for each protein per species. Bin counts were divided by the total number of proteins in each species datasets to yield fractions of evaluated proteins, aiding cross-species comparison. The vertical red line corresponds to the average protein sequence identity.

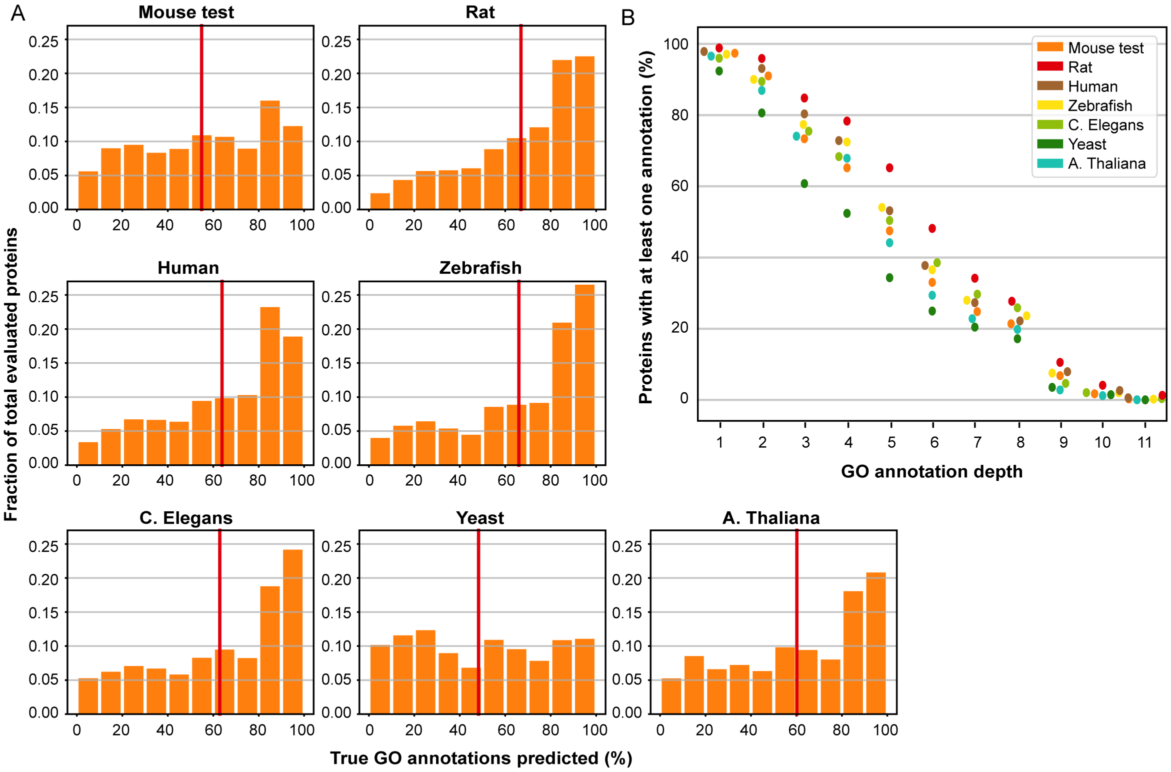


Figure S.7: (A) Distributions of the percentage of predicted real annotations for each protein per species. Real annotations are all the annotations present in the species datasets, including the non-evaluated GO terms. Bin counts were divided by the total number of proteins in each species datasets to yield fractions of evaluated proteins, aiding cross-species comparison. Annotations were predicted by an MLP classifier trained using baseline SeqVec protein-level embeddings using the cross-species datasets. The vertical red line corresponds to the average assigned percentage of true annotations. (B) Percentage of proteins with at least one true positive predicted annotation in relation to the GO term depth. Predictions were made using the same MLP classifier trained using baseline SeqVec protein-level embeddings on the mouse training set. Species are indicated by colour.

| **Species** | **Average IC non evaluated GO terms** |
|---|---|
| Mouse validation | 10.3 (0.2) |
| Mouse test | 10.3 (0.2) |
| Rat | 11.6 (0.3) |
| Human | 12.2 (0.4) |
| Zebrafish | 10.1 (0.3) |
| C. elegans | 10.5 (0.4) |
| Yeast | 10.8 (0.5) |
| A. thaliana | 11.3 (0.8) |

Table S.2: Average Information Content (IC) of the non-evaluated GO terms in protein-centric evaluation per species. Parenthesis indicate standard deviation.

A Biological Process

| Species | # selected proteins | # GO terms | # GO terms evaluated protein-centric (% of terms) | # GO terms evaluated term-centric (% of terms) |
|---|---|---|---|---|
| Mouse | 13.795 | 15.482 | - | - |
| Mouse training | 9.836 | 14.360 | - | - |
| Mouse validation | 1.992 | 9.194 | 8.553 (93%) | 4.335 (47%) |
| Mouse test | 1.967 | 9.124 | 8.528 (93%) | 4.400 (48%) |
| Rat | 6.526 | 13.978 | 12.991 (93%) | 7.023 (50%) |
| Human | 16.067 | 15.434 | 13.984 (91%) | 7.264 (47%) |
| Zebrafish | 2.204 | 5.903 | 5.594 (95%) | 2.710 (46%) |
| C. elegans | 2.888 | 5.127 | 4.725 (92%) | 2.580 (50%) |
| Yeast | 5.122 | 5.192 | 4.139 (80%) | 2.382 (46%) |
| A. thaliana | 11.257 | 5.607 | 4.099 (73%) | 2.432 (43%) |

B Cellular Component

| Species | # selected proteins | # GO terms | # GO terms evaluated protein-centric (% of terms) | # GO terms evaluated term-centric (% of terms) |
|---|---|---|---|---|
| Mouse | 14.263 | 1.932 | - | - |
| Mouse training | 10.144 | 1.850 | - | - |
| Mouse validation | 2.077 | 1.867 | 1.115 (60%) | 585 (31%) |
| Mouse test | 2.042 | 1.815 | 1.143 (63%) | 569 (31%) |
| Rat | 6.655 | 1.695 | 1.633 (96%) | 881 (52%) |
| Human | 17.107 | 1.950 | 1.807 (93%) | 947 (49%) |
| Zebrafish | 2.269 | 879 | 863 (98%) | 443 (50%) |
| C. elegans | 2.944 | 924 | 890 (96%) | 502 (54%) |
| Yeast | 5.416 | 1.043 | 811 (78%) | 467 (45%) |
| A. thaliana | 12.096 | 919 | 743 (81%) | 457 (50%) |

Table S.3: The number of (A) biological process and (B) cellular component GO terms (evaluated) in each species. During protein-centric evaluation GO terms overlapping with the Mouse training set are considered. During term-centric evaluation only GO terms with at least 3 annotated proteins and overlapping with the Mouse training set are considered. The mouse dataset consists of the mouse training, validation and test sets combined.

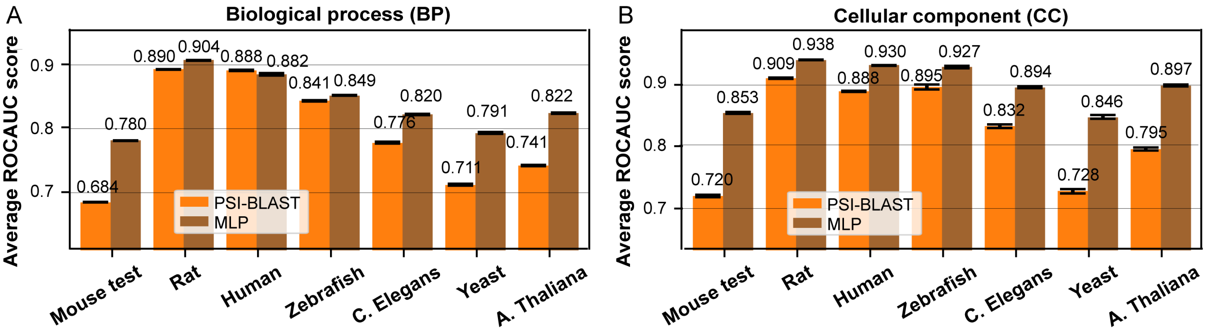


Figure S.8: The average term-centric ROCAUC performance over all the GO terms per species for the MLP classifier (brown) for (A) biological process GO terms and (B) cellular component GO terms. Performance is compared to baseline PSI-BLAST (orange). The coverage C is shown inside the bars. Errorbars denote 95% confidence intervals estimated using 100 bootstraps.

### Materials and Methods

#### Protein-centric metrics

For protein-centric evaluation, we calculated the F1-scores of all test proteins. To calculate the F1-scores, we first calculated for some decision threshold $t \in [0, 1]$, which converted the predicted probabilities into binary class labels, the precision $pr_o(t)$ and recall $rc_o(t)$ for every protein $o$. Using these, we calculated the F1-score $F1_o(t)$ for every protein.

To calculate the average F1-score over all the proteins, we first calculated the average precision and recall over all the proteins for a fixed threshold $t$ using:

$$\text{pr}(t) = \frac{1}{U(t)} \cdot \sum_{o=1}^{U(t)} \text{pr}_o(t) \tag{1}$$

and

$$rc(t) = \frac{1}{V} \cdot \sum_{o=1}^{V} rc_o(t) \quad (2)$$

Note that we calculated the precision only over the $U(t)$ proteins for which at least one prediction was made above threshold $t$ and Recall over all the $V$ proteins. Consequently, we obtained the prediction coverage using:

$$C(t) = \frac{U(t)}{V} \quad (3)$$

which represents the fraction of protein with at least one predicted annotation for threshold $t$, regardless of the prediction being false or true.

Finally, we calculated the average F1-score over all the proteins using:

$$F_1(t) = \frac{2 \cdot \mathrm{pr}(t) \cdot \mathrm{rc}(t)}{\mathrm{pr}(t) + \mathrm{rc}(t)} \quad (4)$$

over all the thresholds $t$. We calculated the maximum $F_1$-score over all thresholds using:

$$F_1 = \max_t \{F_1(t)\} \quad (5)$$

To recreate a real case scenario in the cross-species experiments in which no validation set is available for the test species, we determined the threshold $t$ resulting in the highest $F_{\max}$ score on the mouse validation set. Using this fixed threshold, we calculated the $F_1$-score in the other species.

#### Term-centric metrics

For term-centric evaluation, we calculated the area under the ROC curve (ROCAUC score) of all GO terms. The average ROCAUC score was obtained by averaging over all the evaluated GO terms.
